# Temporal dose inversion properties of adaptive biomolecular circuits

**DOI:** 10.1101/2025.02.10.636967

**Authors:** Eiji Nakamura, Franco Blanchini, Giulia Giordano, Alexander Hoffmann, Elisa Franco

## Abstract

Cells have the capacity to encode and decode information in the temporal features of molecular signals. Many pathways, for example, generate either sustained or pulsatile responses depending on the context, and such diverse temporal behaviors have a profound impact on cell fate. Here we focus on how molecular pathways can convert the temporal features of dynamic signals, in particular how they can convert transient signals into persistent downstream events and vice versa. We refer to this type of behavior as temporal dose inversion, and we demonstrate that it can be achieved through adaptive molecular circuits. Using a cell-free synthetic gene circuit implementing an incoherent feedforward loop (IFFL), we experimentally demonstrate the temporal dose inversion. To understand the design principles of the temporal dose inversion, we analyze adaptive circuit motifs including incoherent feedforward loops (IFFLs) and negative feedback loops (NFLs). Through numerical simulations with expensive parameter exploration, we identify parametric regimes in which these circuits exhibit temporal dose inversion. We further examine more detailed biological models of the IFFL and NFL circuits, including enzymatic signaling models and gene regulatory network models, showing that the IFFL circuits is more likely to exhibit temporal dose inversion compared with the NFL circuits. Finally, we analyze a generalized IFFL topology, and we find that both the time delay in the inhibition pathway and the relative signal intensities of the activation and inhibition signals are key determinants for temporal dose inversion. Together, our results establish a design principle of temporal dose inversion on the adaptive biomolecular circuits and provide mechanistic insight into how molecular networks process temporal information in dynamic signals.

## INTRODUCTION

Biological systems sense and respond to various physiological inputs, such as hormones, cytokines, and growth factors, by processing these signals through complex signaling pathways. These pathways lead to specific biological outcomes not only by producing or eliminating biomolecular components, but also by introducing temporal fluctuations of their concentration or activity^1–4^. A well-known example is the extra-cellular signal-regulated kinase (ERK), which can take different temporal characteristics and accordingly induce different biological outcomes in the mitogen-activated protein kinase (MAPK) network. ERK signals show either a transient or sustained behavior, depending on the input identity^5–8^. Epidermal growth factor (EGF) induces transient ERK activity, which triggers proliferation, whereas nerve growth factor (NGF) induces sustained ERK activity, which is translated to differentiation. ERK also plays an essential role in animal development. In the early Drosophila embryo, a high-dose ERK pulse induces gut endoderm differentiation, while a low-dose ERK pulse triggers cells to adopt neurogenic ectoderm^6^. The importance of temporal features has become evident in many biological contexts, including but not limited to neuronal activity^9^ and hormonal signaling^10–12^, leading to the notion of “temporal signaling” or “temporal coding” by which living organisms encode and interpret information based on the timing, duration, frequency, and amplitude of signals over time.

Here we consider a broad question pertaining to temporal coding: what signaling networks can enable cells to select transient inputs based on their duration and amplitude? It is known that sustained signals and transient signals can be discriminated by simple network motifs such as the incoherent feedforward loop (IFFL) and the negative feedback loop (NFL)^13–18^. The IFFL and NFL are well-known for their adaptive behavior, which means they respond only transiently to changes in their input signal, typically taken to be a sustained amplitude input change. After a transient, the circuit eventually recovers its original state, unless the input perturbs the system beyond its tolerance. Intriguingly, we found that the IFFL and NFL can also produce a sustained output in response to a transient input, indicating their ability to discriminate inputs based on temporal features, and to process them to generate outputs with opposite temporal patterns. Building on these observations, we introduce the notion of temporal dose inversion which occurs if a system produces a sustained output when stimulated with a short-lived input, and vice-versa the system produces a short-lived output in response to a sustained input, as illustrated in Figure 1A and B.

**Figure 1:**
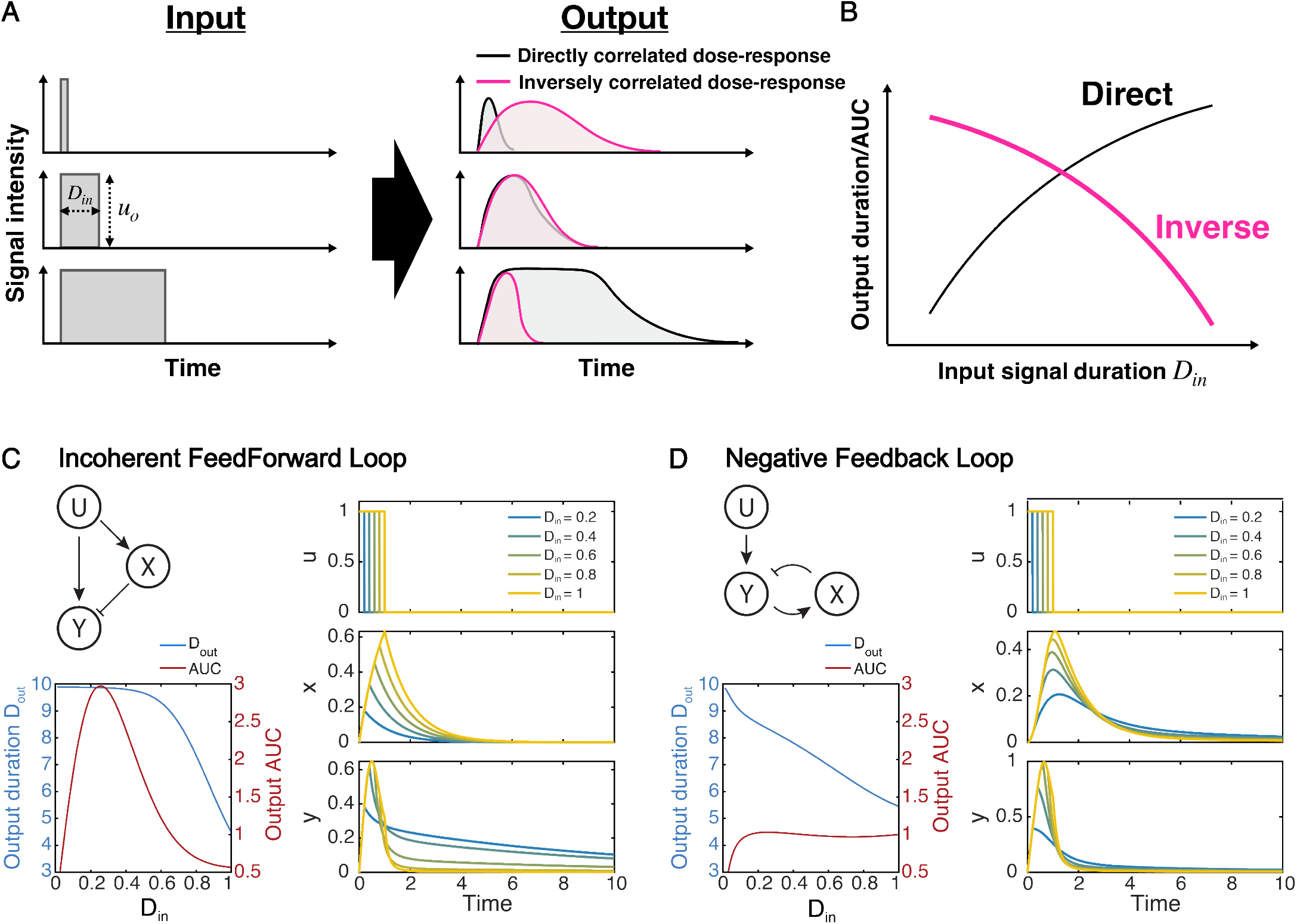
(A) Illustrations of input-output circuit behaviors that maintain or invert the input temporal dose. (B) In a directly correlated dose response, the output dose increases as input dose increases. In an inversely correlated dose response, the output duration decreases as the input dose increases. (C, D) Our simple models for the incoherent feedforward loop (IFFL) and the negative feedback loop (NFL) circuits exhibit temporal dose inversion. Circuit diagrams, and temporal dynamics of (C) IFFL and (D) NFL, responding to pulsatile inputs with varied duration *D*_*in*_ from 0.2 to 1.00, with *a* = 2, *b* = 20, *c* = 0.1, and *p* = 2. Input duration versus output duration or output AUC for (C) IFFL and (D) NFL with c in the range from 0.05 to 1.00 is indicating that IFFL exhibits inverse dose duration (IDD) and inverse dose AUC (IDA) for a certain range of *D*_*in*_, and NFL exhibits IDD in the whole evaluated range of *D*_*in*_.

The IFFL is a versatile motif that features one activation pathway and one inhibition pathway in parallel^17,19–23^. The behavior of the IFFL has typically been studied using step input functions: in this case, the expected response is a transient pulse owing to the presence of a delayed inhibition branch in the circuit^24,25^. The output initially increases and then is repressed, approaching an equilibrium that, in some cases, is identical or sufficiently close to its pre-stimulus state (perfect adaptation)^26^. Under certain assumptions, the output level directly depends on the ratio of the new input level to the pre-stimulus input level (fold-change detection)^27–31^. Furthermore, the IFFL has attracted growing interest as a circuit for processing dynamic signals, especially for detecting and decoding pulsatile signals^13–17^. Some of these studies (computational and experimental) demonstrated that the IFFL circuit or its variants, such as an interlocked IFFL^14,15^, can be utilized as pulse detectors that respond to pulsatile inputs with specific temporal features. Finally, the IFFL can work as a band-pass circuit^14,32,33^.

The NFL is a circuit motif with an inhibitory feedback loop, which is known as a core motif for generating pulses and oscillations in a variety of contexts, including the circadian clock^34^, feedback circuit governing nuclear NF-*κ*B localization^35^, and the p53/Mdm2 network^36^. In addition, under certain conditions the NFL is known for its capability of adaptation and fold-change detection^37–40^, both of which are essential for maintaining the robust operation of cellular functions in noisy environments.

We begin our study by considering simple IFFL and NFL circuit models to investigate the requirements for temporal dose inversion through numerical simulations. Next, we experimentally validate the temporal dose inversion property of an IFFL circuit built using *in vitro* transcriptional gene networks. We then examine whether temporal dose inversion property arises from specific biological mechanisms or emerges purely from network topology by analyzing more complex models meant to capture enzymatic signaling and gene regulatory networks. Finally, we consider a generalized IFFL architecture that is parameterized by the lengths of the activation and inhibition pathways to extract design rules of temporal dose inversion, showing that delayed inhibition is essential for temporal dose inversion. Together, these computational and experimental results provide insights that help us understand how adaptive biomolecular networks decode temporal information, specifically signal duration.

## RESULTS

### 1 IFFL and NFL Models Exhibit Temporal Dose Inversion

#### Reduced order models for IFFL and NFL motifs

Numerous models have been proposed to describe IFFL-based circuits^27,41–43^. Here, we adopt the typical circuit topology of an IFFL composed of an input signal *U*, an intermediate chemical species *X* and an output chemical species *Y* (Figure 1C). We use lowercase letters to indicate concentrations; *u* is the amplitude of input signal *U*, and *x* and *y* represent the concentrations of the species *X* and *Y* respectively. Temporarily denoting the input amplitude by *ū*, we introduce a simple model to describe the IFFL as follows:

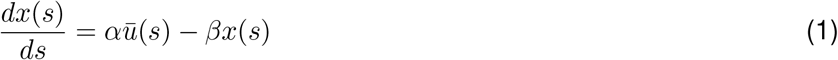

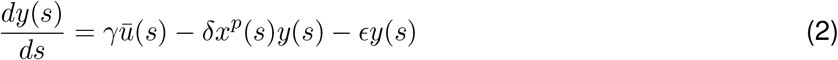

where *α* and *γ* are the input activation rate constants for species *X* and *Y* ; *β* and *ϵ* are self-degradation rate constants for species *X* and *Y* ; δ is the rate constant associated with inhibition of *Y* mediated by *X*, and *p* represents the effective order of the inhibitory reaction, capturing nonlinearities arising from cooperative interactions, multimer formation, or embedded multi-step reaction cascades, while *s* is the time variable. We note that it is essential for our model to be nonlinear, as linear systems do not admit temporal dose inversion (Supplemental information).

To reduce the number of parameters, we rescale time in Equations 1 and 2, obtaining the updated model, which we will analyze in the following without loss of generality:

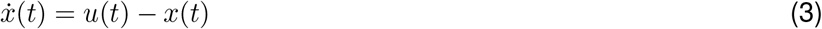

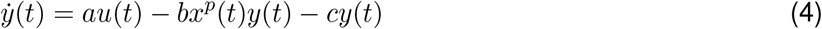

where *t* = *βs, u* = *αū/β, a* = *γ/α, b* = *δ*/*β* and *c* = *ϵ/β*, while 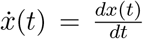 and 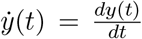. Using Equations 3 and 4, we can evaluate the circuit dynamics with respect to parameters *a, b, c*, and *p*. We examine the response of the model to a rectangular input function of finite duration *D*_*in*_ with amplitude *u*_0_:

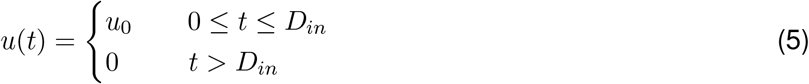

For the NFL, we adopt the simple circuit topology (Figure 1D) and obtain the corresponding equations with rescaled time as follows:

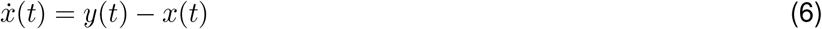

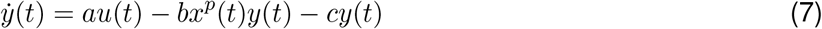

The detailed derivation of Equations 6 and 7 is described in STAR Methods 4. The only difference from the IFFL model is that the first term of Equation 6 is *y*(*t*) instead of *u*(*t*). Throughout numerical simulations using these IFFL and NFL models, the initial conditions are *x*(0) = 0 and *y*(0) = 0. Responses of the simple IFFL model and the simple NFL model for different input durations *D*_*in*_ (from 0.2 to 1.0) are evaluated (Figure 1C and 1D).

#### Measuring temporal inversion properties

To quantify whether the IFFL and NFL have the capacity to invert the temporal features of the input, we measure the output response with two metrics: signal duration and temporal dose. Temporal dose is broadly intended as the area under the curve (AUC, i.e. the integral) of a transient signal, which depends on both its amplitude and its duration. Accordingly, we define two types of temporal dose inversion properties: inverse dose duration (IDD) and inverse dose AUC (IDA). An IDD circuit converts a short pulse into a longer pulse and vice versa, functioning as a duration inverter. An IDA circuit converts a small-AUC input into a large-AUC output and vice versa, functioning as a temporal dose amplifier. When a circuit specifically detects a transient signal, it is expected that the circuit shows the temporal dose inversion properties. Through computational methods, we characterize the temporal dose inversion properties of IFFL and NFL circuits by considering pulsatile inputs of varying duration, and derive parametric and topological conditions for implementing inversion properties.

The IFFL exhibits a long-lasting response to a shorter input duration (e.g. *D*_*in*_ = 0.2), and thus shows the capacity for IDD (Figure 1C). To quantitatively evaluate its temporal dose inversion properties, we computed the output duration *D*_*out*_ and the area under the curve (*AUC*). *D*_*out*_ is calculated as the square root of the variance in the time domain, where the variance is derived as the difference between the second moment 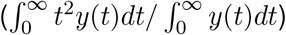 and the square of the mean 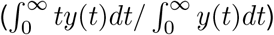. This approach is analogous to the calculation of the standard deviation in probability distributions. *AUC* is the integration of the output signal *y*(*t*) from *t* = 0 to *t* = 80, as described in detail in STAR Methods 4. *D*_*out*_ and *AUC* are evaluated as a function of *D*_*in*_ (lower left panels of Figure 1C and D). *D*_*out*_ monotonically decreases as *D*_*in*_ increases: *D*_*out*_ and *D*_*in*_ are inversely correlated, and therefore the system has the inverse dose duration (IDD) property. Conversely, *AUC* increases when 0 *< D*_*in*_ *<* 0.25, has a peak at *D*_*in*_ = 0.25, and decreases when *D*_*in*_ *>* 0.25. Hence, we argue that the IFFL exhibits the inverse dose AUC (IDA) property just in a range of *D*_*in*_ values (*D*_*in*_ *>* 0.25). The NFL circuit also exhibits IDD, as *D*_*out*_ is monotonically decreasing with respect to *D*_*in*_, while it does not exhibit IDA, since *AUC* increases for increasing *D*_*in*_ (Figure 1D). We will compare the likelihood of the IDA property for IFFL and NFL in Section 3.

#### Parametric regimes enabling temporal dose inversion

To understand how each parameter shapes the circuit dynamics and temporal dose inversion, we first identify a set of nominal parameter values that exhibit temporal dose inversion through trial and error. We then vary *a, b* and *c* individually by one order of magnitude around their nominal values, taking input durations *D*_*in*_ = 0.1 and *D*_*in*_ = 1 (Figure 6). The activation parameter *a* mostly changes the amplitude of the output, without significantly affecting its duration. The inhibition parameter *b* and the degradation parameter *c* change the output response more drastically. As *b* or *c* increases, the output duration decreases, because a larger value of *b* or *c* brings the output level more quickly back to its basal level. However, when *D*_*in*_ = 1, the output duration does not perceptibly decrease as *c* increases; this happens because, owing to the longer input duration, the effect of the inhibition term *bx*^*p*^*y* is dominant.

Heat maps of *D*_*out*_ and *AUC* displayed in the *D*_*in*_-*a, D*_*in*_-*b* and *D*_*in*_-*c* planes reveal the emergence of temporal dose inversion and the influence of each parameter (Figure 2). When there is a cross-section at a fixed parameter value along which *D*_*out*_ decreases over a range of *D*_*in*_ (e.g., *b* = 20 in the left middle panel of Figure 2B), the circuit exhibits the IDD property for that parameter set. Similarly, a decrease in *AUC* at a cross-section indicates the IDA property.

**Figure 2:**
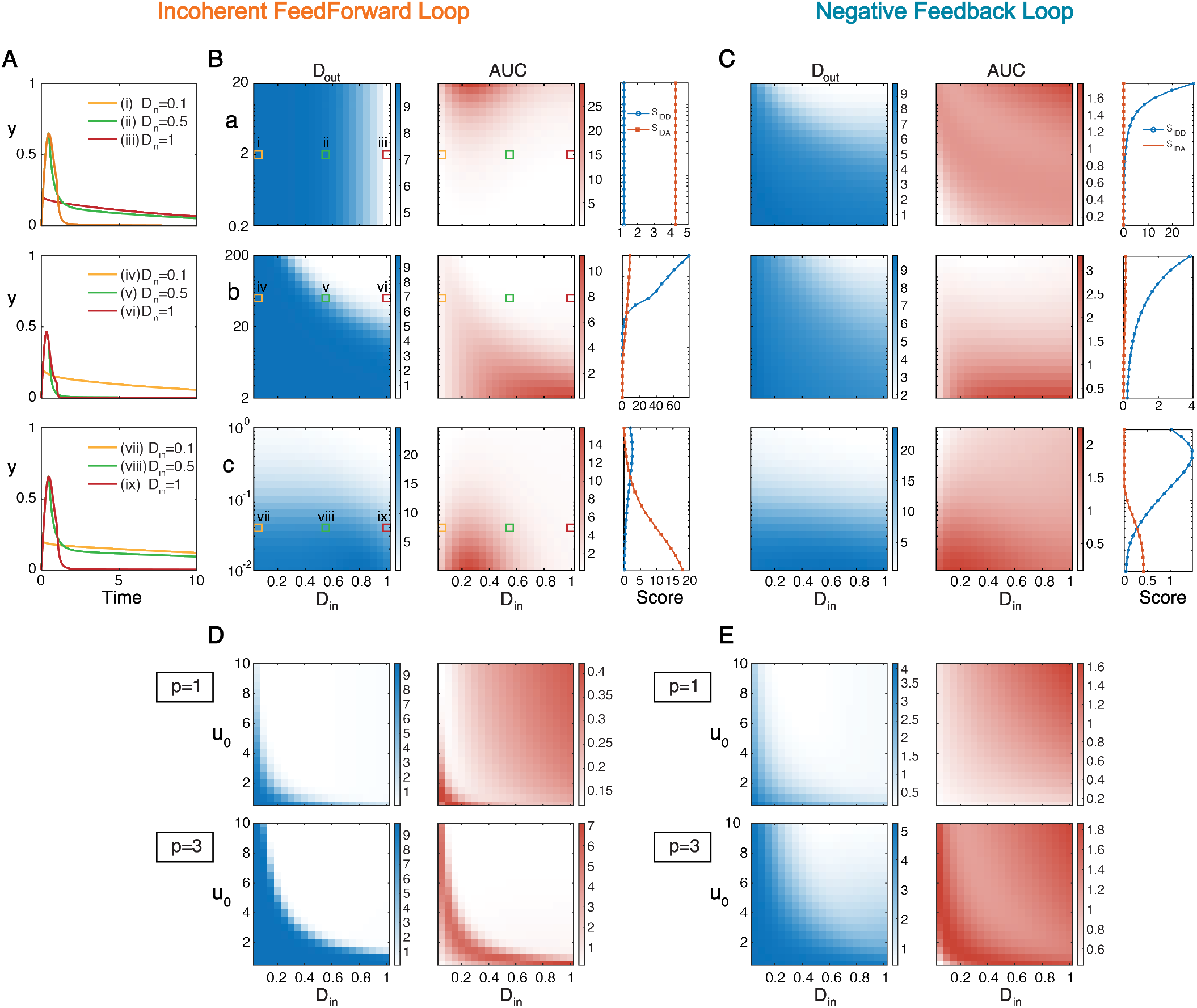
Evaluating the inverted dose-response as a function of individual parameters related to output production (*a*), inhibition (*b*), and degradation (*c*). (A) Time evolution of the output *y* with *D*_*in*_ = 0.1, *D*_*in*_ = 0.5, *D*_*in*_ = 1. Each output curve corresponds to the data point indicated with the same Roman numeral in the heat maps in panel B. (B,C) Heat maps showing the values of *D*_*out*_ (blue heat maps) and *AUC* (red heat maps) for the outputs of the IFFL (B) and the NFL (C) in the *D*_*in*_-*a, D*_*in*_-*b, D*_*in*_-*c* planes respectively. Here, *D*_*in*_ is linearly varied from 0.1 to 1 in steps of 0.05. In the same row or the same column, heat maps have the same axes units (only shown in the leftmost or bottom panels). Each row shows heat maps for a different parameter: *a*, top; *b*, center; *c*, bottom.

To systematically assess the IDD and the IDA property, we calculate an IDD score (*S*_*IDD*_) and an IDA score (*S*_*IDA*_) as follows. Given an input duration interval *D*_*in*,1_ *< D*_*in*_ *< D*_*in*,2_ for which the function *D*_*out*_(*D*_*in*_) has a negative derivative,

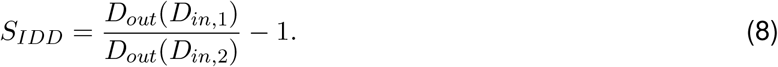

If there are multiple input duration intervals in which the function *D*_*out*_(*D*_*in*_) has a negative derivative, we choose our score *S*_*IDD*_ as the maximum value of 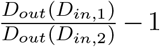. If the function *D*_*out*_(*D*_*in*_) is monotonically increasing or constant, *S*_*IDD*_ = 0, which means that no IDD property is observed in the considered parameter set. *S*_*IDA*_ is computed analogously for the function *AUC*(*D*_*in*_). By computing *S*_*IDD*_ and *S*_*IDA*_ values for each parameter set, we can quantify to which degree the IDD and IDA properties hold within that parameter set. More details on the mathematical computation of *S*_*IDD*_ and *S*_*IDA*_ are provided in STAR Methods 18.

The IFFL shows both the IDD and IDA properties within a certain range of each parameter (Figure 2A and B). While the activation parameter *a* does not change *D*_*out*_ (and therefore does not affect the IDD property), a higher value of *a* favors an apparent IDA property. The inhibition parameter *b* and the degradation parameter *c* significantly affect both *D*_*out*_ and *AUC*; as *b* or *c* increases, the circuit dynamics accelerates, which can be seen in the fact that the region with large *D*_*out*_ expands as *b* or *c* decreases. The dependence of *AUC* on *b* is more complicated than that of *D*_*out*_. Although *AUC* increases as *b* decreases because of a smaller inhibition effect, decreasing *b* leads to the loss of the IDA property: for instance, along a cross-section with *b* = 2 (Figure 2B middle), *AUC* monotonically increases as *D*_*in*_ increases (hence, there is no IDA property). Therefore, *b* should not be too small to obtain the IDA property at least for a range of *D*_*in*_ values, whereas a larger value of *a* and lower value of *c* are preferable to ensure the IDA property at least in some ranges of *D*_*in*_.

The NFL can also exhibit the IDD and IDA properties, but for a smaller set of parameters with respect to the IFFL (Figure 2C). Concerning the IDD property, differently from the IFFL circuit, the NFL shows a strong dependence on the activation parameter *a*; in fact, the inhibitory feedback from *X* has to be triggered by *Y* and *a* determines how quickly *Y* activates *X*, which eventually affects *D*_*out*_. Concerning the IDA property, it is worth pointing out that the color scales of the heat maps in Figure 2C (right) are smaller than those in Figure 2B (right), suggesting that, in the NFL, it is more difficult to identify a cross-section at a parameter value along which we see an apparent IDA property. Here is an explanation as to why the IFFL is more likely to exhibit the IDA property. In the IFFL, the activation and inhibition pathways can be tuned independently, so we can choose the parameters so that the output node *Y* is immediately activated, while *X* is activated with a certain (tunable) delay. Therefore, if *D*_*in*_ is smaller than a threshold, input *U* only activates *Y*, without activating the inhibition signal from *X*. Once *D*_*in*_ exceeds the threshold, both the activation and inhibition pathways are triggered, which eventually decreases *AUC*. This is the mechanism that allows us to attain the IDA property in the IFFL. In contrast, in the NFL, the activation pathway and the inhibition pathways acting on *Y* are not independent, and are stimulated simultaneously in most cases; in fact, it is relatively difficult to find a condition in which only *Y* is activated, keeping the inhibition feedback from *X* not significantly stimulated, because active *Y* stimulates *X*. This does not mean that the NFL does not exhibit the IDA property for any choice of the parameters, but provides us with a qualitative understanding of why IFFL is more likely than NFL to exhibit the IDA property.

Parameter *p*, the order of the inhibition term, also affects the IDD and IDA properties. *D*_*out*_ and *AUC* are shown as heat maps in the *D*_*in*_-*u*_0_ plane (Figure 2D and E). For both IFFL and NFL, the IDA property is reinforced as *p* increases, namely, the ranges of *D*_*in*_ and *u*_0_ where an apparent inverse relationship occurs and the corresponding dynamic ranges of the *AUC* expand, while the IDD property is not apparently reinforced for both circuits. When *p* = 3, both the IFFL and NFL do not exhibit the IDD and the IDA properties for small *u*_0_ (such as *u*_0_ = 0.5), while they do exhibit the properties for larger *u*_0_. This happens because, when *x* < 1, the inhibition term *bx*^*p*^*y* decreases as *p* increases.

Varying the input amplitude *u*_0_ can also lead to temporal dose inversion (Figure 2D and E). This can be explained by noting that the *x* variable of the IFFL responds to an input (starting from the initial condition *x*(0) = 0) by increasing monotonically and asymptotically reaching its equilibrium *x*_*eq*_ = *u*_0_. Therefore, if *u*_0_ *<* 1, then it is always *x*(*t*) *<* 1, which reduces the effect of the inhibition term *bx*^*p*^*y*. However, if *u*_0_ *>* 1, the degree *p* of the inhibition term reinforces the IDD and IDA properties. Interestingly, all heat maps are symmetric with respect to the diagonal line in the *D*_*in*_-*u*_0_ plot (which is not the bisector, since the two axes have different scales).

## 2 Cell-free demonstration of Temporal Dose Inversion

We experimentally demonstrate that temporal dose inversion occurs in a synthetic IFFL circuit built using cell-free transcriptional components known as “genelets”^21,44,45^.

### Transcriptional genelet circuits

A genelet is a linear, double-stranded DNA construct that contains a T7 promoter with a nick in its template strand (Figure 3A). Transcription from a genelet depends on whether an activator strand (dA) is bound (genelet transcription on) or not (genelet off) to the promoter region, which operates as an input domain. The output of a genelet is its RNA transcript, whose sequence content is completely decoupled from the input. Thus, a genelet can be graphically represented as a node characterized by its independent inputs and outputs (Figure 3A, bottom). In this system, transcription is achieved using bacteriophage T7 RNA polymerase, and RNA is degraded by introducing RNase H^44^, which targets RNA in DNA-RNA complexes. We measure the on/off state of a genelet by labeling genelet templates with a fluorophore (labeled F in Figure 3A), and DNA activators with a quencher (Q in Figure 3A).

**Figure 3:**
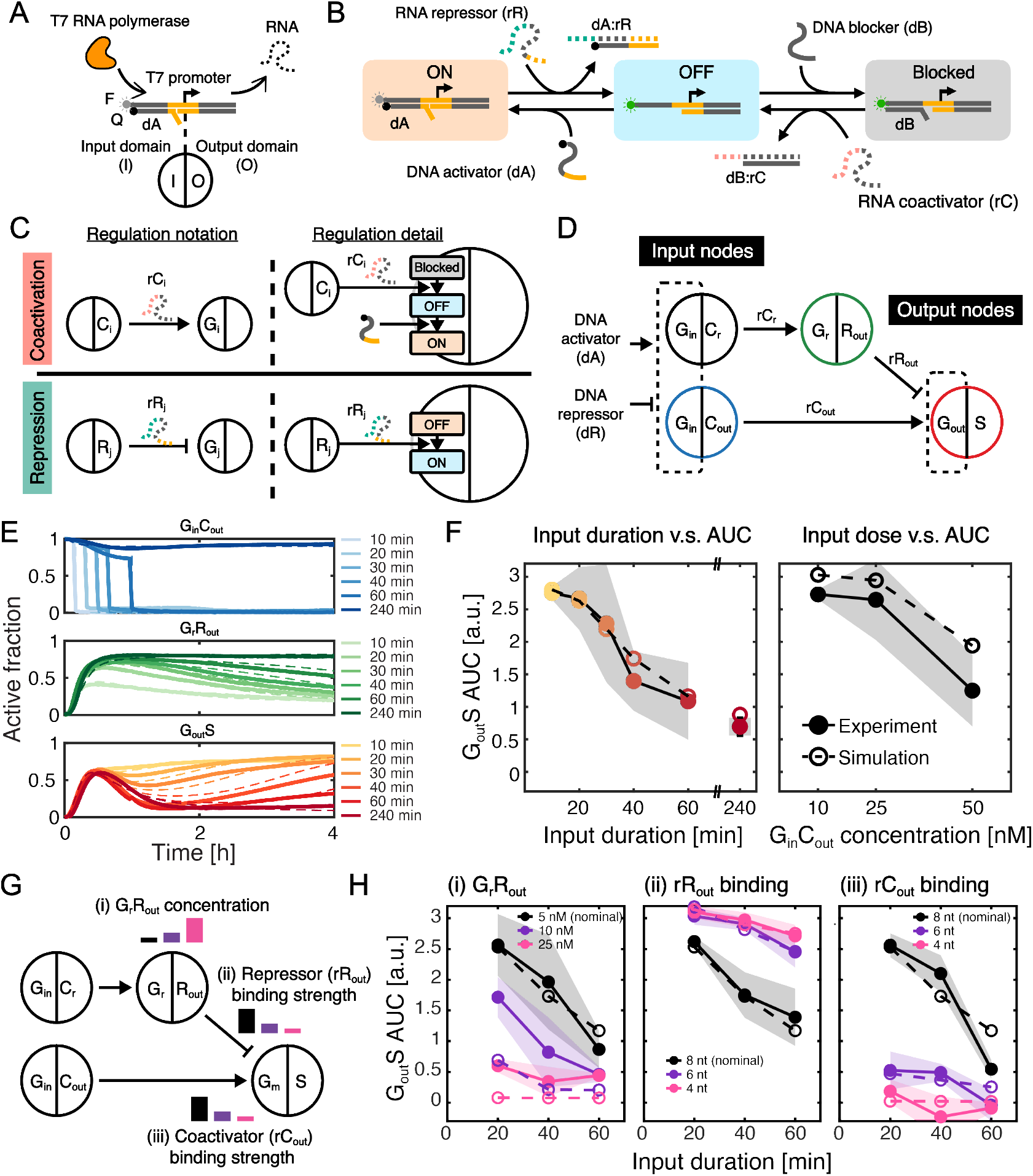
A genelet-based incoherent feedforward loop (IFFL) exhibits the inverse dose-AUC (IDA) property. (A) Schematic of a genelet, in which the T7 promoter (domain in yellow) has a nick on the template strand. The non-template strand of the genelet has a fluorophore (F) on its 5’ end, and dA has a quencher (Q) on its 3’ end. (B) Genelet regulation through toehold-mediated strand displacement. The DNA activator (dA) has a toehold domain that enables its displacement by an RNA repressor (rR), repressing transcription from the genelet. Transcription can also be prevented in the presence of a DNA blocker (dB) due to the incomplete T7 promoter. The DNA blocker also has a toehold domain, which enables its removal by an RNA coactivator (rC). (C) We represent genelet networks with nodes (genelets) interconnected by arcs (regulatory interactions). Each genelet node is depicted by a circle separated into two parts: the left half shows the input domain, and the right half shows the output domain. Arcs are represented with the symbol “→” when associated with an RNA coactivator, and with “ ⊣” if the associated signal is a repressor. (D) Schematic of our genelet-based incoherent feedforward loop (IFFL). The input is received by two separate genelets (G_in_C_out_ and G_in_C_r_) with the same activation domain. Transcription from the input nodes is turned on by a single DNA activator (dA_in_), and repressed by a single DNA repressor (dR_in_). Not shown in this schematic is one additional genelet, G1I1, at very low concentration (1 nM), which suppresses the transcription leakage of G_r_R_out_. Genelets meant to be activated by RNA coactivators are initially in blocked states. A fully-detailed schematic of the system is provided in the Supplemental Information (Figure4). (E) Responses of the genelet IFFL to varying input durations. The three graphs display normalized active levels of the three genelets: G_in_C_out_ (top), G_r_R_out_ (middle), and G_out_S (bottom) responding to to six different input durations (10, 20, 30, 40, 60, and 240 min). Each plot shows the result of a single replicate. Additional replicates are shown in the Supplemental Figure 12. G_in_C_out_ and G_in_C_r_ are initially in ON state, and T7 RNA polymerase is added at *t* = 0 to initiate the circuit operation. The active levels of G_in_C_out_ and G_in_C_r_ are shaped into pulses by adding dR_in_ at specific time points (indicated by “↓”), which determine the pulse durations (only G_in_C_out_ is monitored). (F) Areas under the curves of the circuit responses depend on the input durations and input node concentrations. *AUC* of G_out_S versus input duration (top) and AUC of G_out_S versus input node concentrations (bottom). Panel F (top) was generated from the trajectories shown in (E) and additional replicates shown in the Supplemental Information, while Panel F (bottom) was generated from the trajectories shown in the Supplemental Figure 12 and 13. Throughout the experiments we maintained a ratio of [G_in_C_out_] to [G_in_C_r_] equal to five. Thus, in Panel F (bottom), [G_in_C_r_] also varies accordingly. For each data point, we visualize the mean ± standard deviation (*n* = 3). (G, H) The IDA behavior of the circuit was modulated by varying (i) [*G*_*r*_ *R*_*out*_], (ii) the binding rate of *rR*_*out*_, and (iii) the binding rate of *rC*_*out*_. (G) shows a schematic indicating where these parameters act in the circuit. (H) shows *AUC* of G_out_S versus input duration.

Binding of an activator to its input domain is competitively regulated by the presence of a DNA blocker (dB). Activator and blocker DNA strands can themselves be sequestered by external regulatory RNAs, an RNA co-activator (rC) and an RNA repressor (rR), to provide an additional layer of regulation (Figure 3B). Genelets can easily be interconnected in a cascade, where the genelet upstream can transcribe a coactivator or a repressor RNA for the downstream genelet (Figure 3C). Furthermore, because input domains and regulatory-RNA sequences can be freely recombined, it is possible to realize nearly arbitrary network topologies. These networks can be represented as graphs where nodes are associated with genelets, and arcs with the RNA input/output molecules functionally interconnecting the genelets^21^ (Figure 3C,D).

### A genelet-based IFFL

Figure 3D shows our IFFL circuit, which is consistent with previous work by Schaffter and coauthors^21^. Two input genelets, G_in_C_out_ and G_in_C_r_, share an identical input domain; thus a single DNA activator (dA_in_) activates both input genelets, and a single DNA repressor (dR_in_) represses both of them. The circuit implements an IFFL, in which one pathway directly activates the output genelet (G_in_C_out_ →G_out_S), while the other pathway indirectly inhibits the output node through the intermediate node (G_in_C_r_ →G_r_R_out_ ⊣G_out_S). We tuned the dynamics of the circuit to achieve a desired operating point by varying the concentrations of the individual genelets and the enzymes T7 RNA polymerase and RNase H. All working concentrations used in our experiments are listed in the Supplemental Information. These values were established through a set of preliminary experiments so that the timescale of experimentally observed on/off transitions of input, intermediate, and output genelets (measured through fluorescence spectroscopy) would be consistent with the timescale of computational simulations discussed in the previous sections. For a detailed discussion of how genelet and enzyme concentrations shape circuit dynamics, we refer the reader to the work by Schaffter and coauthors^21^.

### Experimental Validation of Temporal Dose Inversion

We generated a finite-duration stimulus for the genelets by manually adding DNA activator and a complementary DNA repressor for the input genelets. The repressor displaces the activator due to the presence of a toehold, and turns the input genelets off, a reaction that equilibrates within minutes (Figure 3E, top). Thus, the time at which the DNA repressor is added determines the input duration, which we define as the time interval during which the two input genelets (G_in_C_out_ and G_in_C_r_) remain active. We measured the responses of the IFFL genelets to varying input durations (10, 20, 30, 40, 60, and 240 min). In Figure 3E we report the corresponding active fraction of one input genelet, of the intermediate repressor genelet, and of the output for the average of three experimental replicate (all the individual replicates are shown in Figure 12). We found that the active fraction of the intermediate repressor gene G_r_R_out_ increased with longer input pulses, and persisted after the inputs were repressed. Consequently, the response of the output node G_out_S evolved from sustained to transient as the input duration increased. (A slight decay in the active fraction of G_in_C_out_ during this window arises from genelet auto-inhibition, as reported previously^21^).

In this experimental validation, we focus on the evaluation of the inverse dose-AUC (IDA) property, as *AUC* provides a more biologically relevant metric. To quantify the IDA property of the circuit, we evaluated the *AUC* of G_out_S response to varying input durations and input node concentrations, as shown in Figure 3F. The *AUC* of G_out_S (Figure 3F) was inversely correlated with both input duration and input node concentration, indicating that the genelet-based IFFL exhibits the IDA property.

The sustained activation of the output node in response to short input durations arises from the accumulation of a substantial amount of *rC*_*out*_ prior to the shutdown of the input nodes. This accumulated *rC*_*out*_ remains sufficient to sustain output activation over the timescale of the experiment, even in the presence of RNase H. In contrast, repression by *rR*_*out*_ is delayed because it requires the activation of *G*_*r*_*R*_*out*_ and the accumulation of *rR*_*out*_ to a level sufficient to effectively repress the output node. Consequently, the circuit exhibits a regime in which activation persists before repression becomes dominant.

Around the transition from sustained to transient output (30 - 40 min), the standard deviation (indicated by the error bar) among three replicates was larger, probably reflecting increased sensitivity to experimental variations of concentration and activity of T7 RNA polymerase and RNase H, which is well-documented in the literature^45^. We also computationally confirmed that variations in enzymatic parameters affect the behavior of our circuit (the Supplemental Figure 14). Overall, these experiments confirm that a genelet-based IFFL can realize the IDA property *in vitro*.

### IDA critically depends on the speed and strength of activation and inhibition

We experimentally characterized how key parameters affect the IDA property (Figure 3G and H). We varied three conditions: (i) the total concentration of *G*_*in*_*C*_*r*_, which determines the speed of the inhibition pathway; (ii) the toehold length of *dA*_*out*_, which determines the binding strength between *rR*_*out*_ and *dA*_*out*_ (inhibition), and (iii) the toehold length of *dB*_*out*_, which determines the binding strength between *rC*_*out*_ and *dB*_*out*_ (activation). Varying these conditions enables us to explore how temporal dose inversion depends on the relative speed and strength of activation and inhibition pathways. These individual parameter variations monotonically changed *AUC*, which eventually eliminated the IDA property, showing that balanced activation and inhibition strength are essential.

A complete mathematical model of the genelet-based IFFL (Supplemental Information), fitted to the data, generally reproduces experimental observations (dashed lines in Figure 3 E, F, H). The model includes 32 variables, indicating that while the system’s behavior is predictable, it is also much more complex than what is depicted in the schematic.

## 3 Temporal Dose Inversion In Signaling and Regulatory Networks

We next ask whether temporal dose inversion arises from specific biological mechanisms or instead solely from network topology. We consider more detailed IFFL and NFL models based on enzymatic signaling and gene regulatory networks (GRNs). In the enzymatic signaling models, the input *U* represents a transient upstream signal or kinase activity, whereas in gene regulatory networks it represents transient transcription factor availability. These two models capture distinct biological mechanisms of activation and inhibition: direct post-translational regulation versus transcriptional regulation. Comparing these two models allows us to test whether temporal dose inversion depends on specific biochemical implementation or instead arise from the underlying IFFL topology. Although real signaling dynamics are not strictly rectangular, approximating transient inputs as finite-duration pulses is effective in capturing temporal dose inversion, which depends primarily on nonlinear dynamics and time-integrated features rather than the precise input shape.

Our signaling model for the IFFL is based on the work by Gerardin et al.^46^:

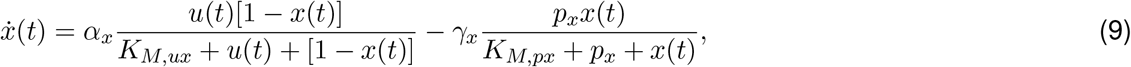

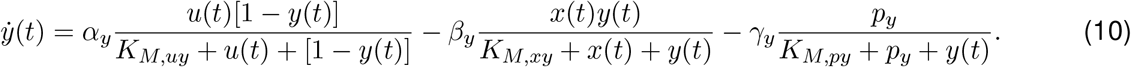

All reactions assume enzymatic signaling and involve two species, *X* and *Y*, and one input, *U*. Each species is either in an active or inactive state. Activation and deactivation are scaled by reaction parameters *α* and *β*; *K*_*M*_ parameters are Michaelis-Menten constants; *p*_*x*_ and *p*_*y*_ are the concentrations of two basal deactivators, while *γ*_*x*_ and *γ*_*y*_ are the corresponding deactivation rate constants. Subscripts indicate the species associated with a particular parameter: for example, *α*_*x*_ is the activation rate constant of *X* and *K*_*M,xy*_ is the Michaelis-Menten constant of enzyme *X* acting on substrate *Y*. To simplify the model, we assume that the total concentration of each species is conserved and normalized to 1. Further, we assume total quasi-steady-state approximation (tQSSA), which is valid for a wider range of parameter spaces than the conventional QSSA^47^.

The gene regulatory network (GRN) model of the IFFL is as follows:

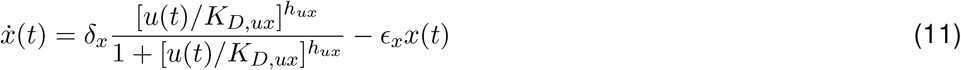

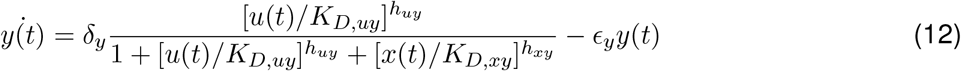

where *x* and *y* denote concentrations of proteins (transcription factors). Parameters δ are maximal activation rates, while rate constants *ϵ* are associated with protein dilution/degradation. *K*_*D*_ is the dissociation constant and *h* is the Hill coefficient of a transcription factor from its target promoter.

The NFL models based on enzymatic signaling and gene regulatory networks are reported in the STAR Methods 4.

We evaluate the likelihood of the IDD or IDA properties of these four models in response to a rectangular input function as in Equation 5, with a constant amplitude (*u*_0_ = 0.01 for the signaling model, and *u*_0_ = 1 for the GRN model), and varying duration *D*_*in*_, assuming zero initial conditions. We examine 10000 parameter sets randomly sampled from a log uniform distribution (STAR Methods 4, reported in Table 4), and we count the sets for which the system exhibits IDD or IDA.

We find that the emergence of temporal dose inversion depends on both biological mechanism and network topology. Signaling models exhibit higher IDD and IDA scores when compared to the GRN models, suggesting a strong dependence on mechanism (Figure 4). At the same time, the IDA property is consistently observed more frequently in the IFFL than in the NFL across both model classes, indicating a dependence on topology. The signaling IFFL model exhibits a particularly high likelihood of achieving large IDA scores, likely due to the independent activation and inhibition pathways inherent to the IFFL motif. Note that a large fraction of parameter sets with positive IDD scores does not necessarily imply robust IDD behavior, as strong IDD (e.g., *S*_*IDD*_ *>* 0.1) is less prevalent and depends on the range of input durations. Therefore, these fractions should not directly be interpreted as a probability of robust temporal dose inversion.

**Figure 4:**
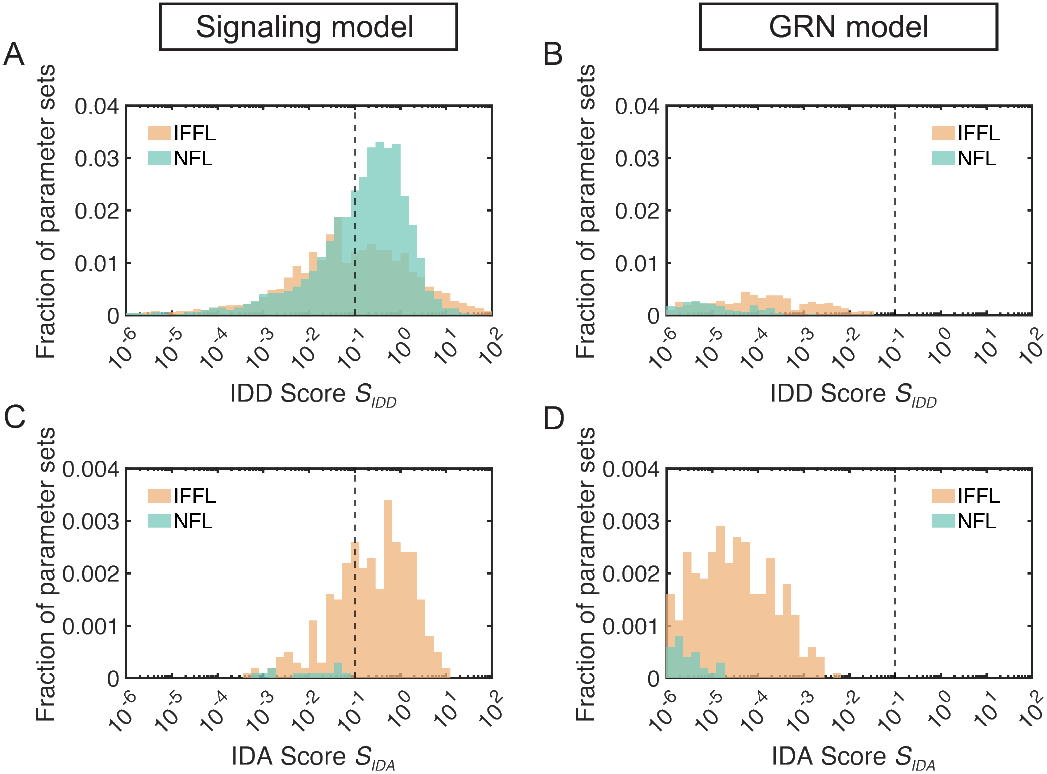
The IDA property is more likely to emerge in the IFFL motif. Histograms of IDD and IDA scores for 10000 randomly-chosen parameter sets. Each parameter is chosen from a log-uniform distribution in the interval specified in STAR Methods 4. For the IFFL signaling model, (A) IDD score histogram and (C) IDA score histogram. For the NFL signaling model, (B) IDD score histogram and (C) IDA score histogram. (E, F) IDD score histograms of gene regulatory model of the (E) IFFL and (F) NFL; IDA scores for these circuits are not shown because a negligible number of parameter sets present a positive score.

IDD and IDA scores of the GRN models are consistently low (below 0.1, marked as a dashed line) when compared to signaling models. A possible explanation is that the GRN inhibitory interaction reduces the production rate of the output species, rather than eliminating the output species; in addition, the output decay does not depend on *X*. Another possible reason is that our model does not capture delays that may occur during translation. If we allow additional intermediate nodes to introduce additional time delay in the GRN model, the IDD and IDA properties can be achieved (Figure S 9).These observations suggest that, within the simplified GRN framework considered here, transcriptional gene regulation alone is less likely to realize temporal dose inversion without additional protein–protein interactions or explicit time delays.

In the Supplement, Figure 15, we explore two additional signaling circuits, one that combines the IFFL and NFL topology (hereinafter, referred to as IFFL+NFL), and one that includes an antithetic integral feedback controller^40,48^. The IFFL+NFL circuit exhibits an intermediate behavior between the IFFL and NFL, suggesting that combining these two motifs does not produce a synergistic effect in terms of temporal dose inversion. The circuit with the antithetic integral feedback controller exhibits temporal dose inversion with a frequency comparable to that of the IFFL. Investigating the relationship between temporal dose inversion and adaptation in circuits with antithetic integral feedback would be an interesting direction for future work, although it is beyond the scope of the present study.

## 4 Signal Amplification and Time Delay Enable Temporal Dose Inversion in Generalized IFFLs

Given that IFFL models show the IDA property more frequently than NFL models, we examine generalized IFFL architectures with varying pathway lengths to assess whether temporal dose inversion depends on topological parameters, seeking to extract design principles to achieve IDA (the analysis can be readily extended to IDD using the same methods). Our generalized IFFL model is:

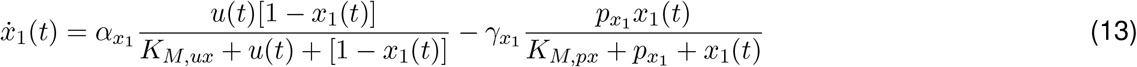

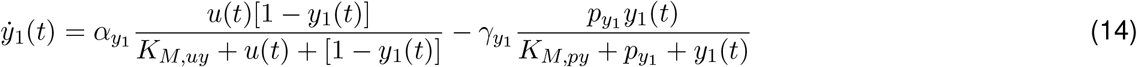

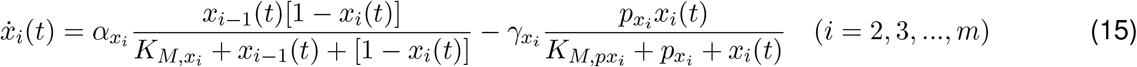

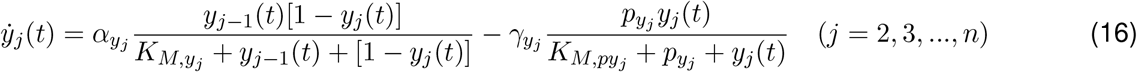

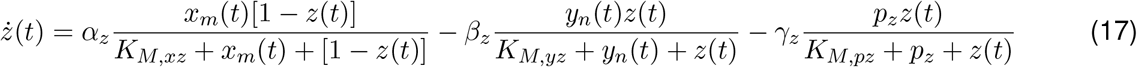

Parameters are labeled consistently with the IFFL signaling model in the previous section, and we introduce a variable pathway length defined by *n* and *m*. Species *X*_*i*_ (*i* = 1, …, *m*) are in the activation pathway, and *Y*_*i*_ (*i* = 1, …, *n*) are in the inhibition pathway. Node *Z* is the output node, activated by *X*_*m*_ and inhibited by *Y*_*n*_. To simplify the analysis, we assume 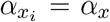 for all *i*, and 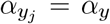 for all *j*. The initial conditions of all variables are set to zero.

We investigate the dependence of IDA on the pathway length by varying *m, n*) and pathway strength by changing *α*_*x*_, *α*_*y*_, *α*_*z*_ and *β*_*z*_ (Figure 5).

**Figure 5:**
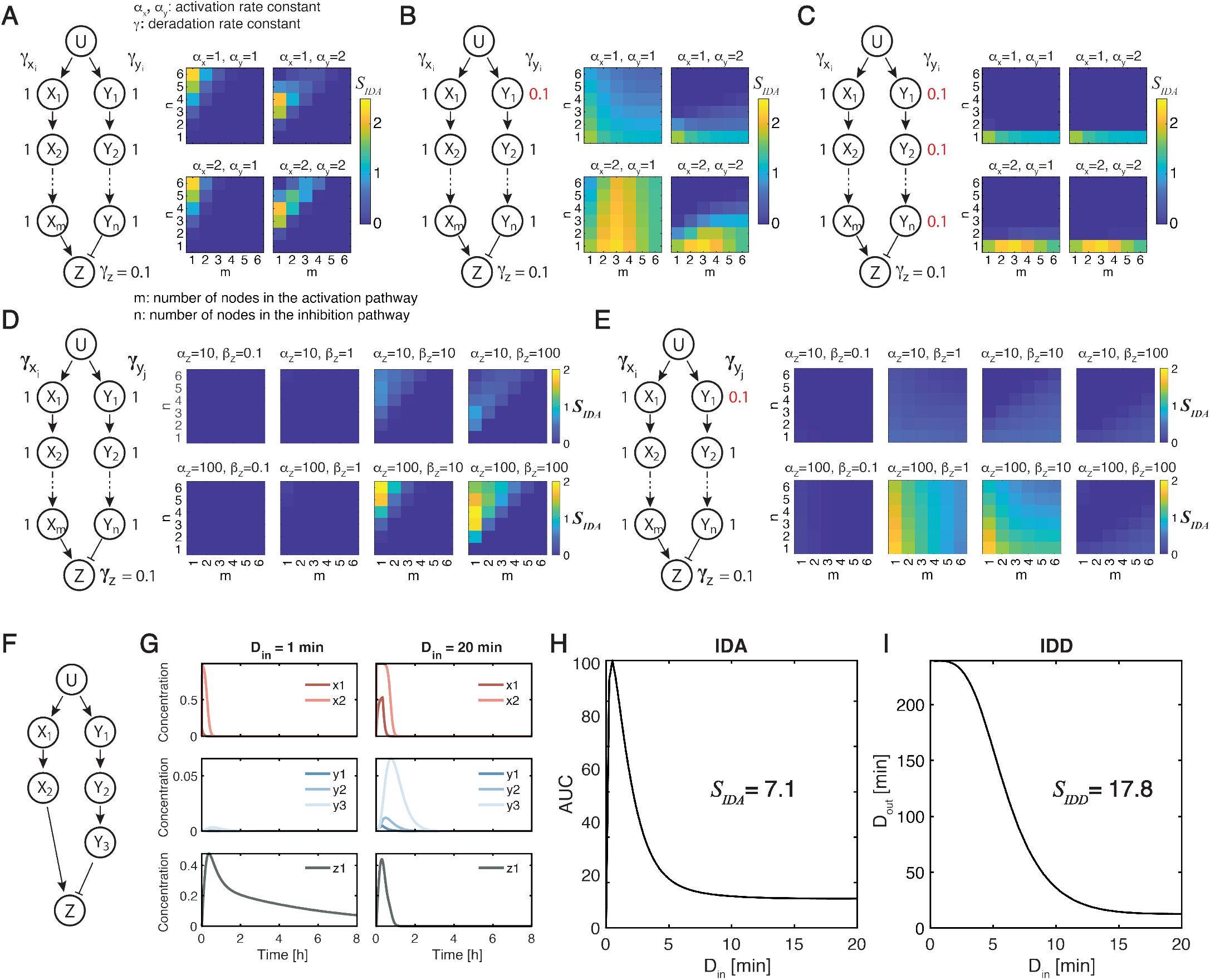
Generalized IFFL circuits show that temporal dose inversion depends on the relative strength of activation and inhibition pathways. (A to C) Each panel shows the circuit diagram of a generalized IFFL network, along with heat maps of IDA scores *S*_*IDA*_ in the *m*-*n* plane for different values of *α*_*x*_ and *α*_*y*_, reported above each heat map. The basal deactivation rates are reported next to each node in the circuit diagram: in all three cases, 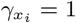 for all *i* = 1, …, *m*, and moreover (A) 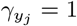 for all *j* = 1, …, *n*; (B) 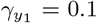, and 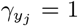 for *j* = 2, …, *n*; (C) 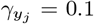 for all *j* = 1, …, *n*. Other parameters are kept constant at their nominal values listed in Table 1. The input amplitude is set at *u*_0_ = 0.1; *D*_*in*_ is logarithmically varied from *D*_*in*_ = 0.1 to *D*_*in*_ = 10. (D, E) Similarly, heat maps of IDA scores *S*_*IDA*_ in the *m*-*n* plane are shown for different values of *α*_*z*_ and *β*, reported above each heat map. (F to I) An example of a circuit with a remarkably high IDA score. (F) Circuit diagram of a generalized IFFL network with *m* = 2 and *n* = 3. (G to I) Time evolution of the output responses of the circuit shown in (F) to input pulses of *u*_0_ = 0.005 with two different input durations: *D*_*in*_ = 1 min (left), and *D*_*in*_ = 20 min (right). Parameter values are listed in Table 3. (H,I) Plots of *AUC* as a function of *D*_*in*_ (H) and *D*_*out*_ (I) as a function of *D*_*in*_ obtained from numerical simulations for the circuit shown in (F). *D*_*in*_ was linearly increased from 0 to 20 minutes with steps of 0.25 min. Note that *D*_*out*_(*D*_*in*_ = 0) is not included in the plot (G) because *D*_*in*_ = 0 gives *y*(*t*) = 0, with which *D*_*out*_ cannot be calculated by Equation 42. When not varied, the parameters are kept constant at their nominal values listed in Table 1, 2 and 3

First, we fix *α*_*z*_ and *β*_*z*_ and consider three different parameter sets of 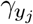 (*j* ∈ {1, 2, …, *n*}) (Figure 5A to C). For 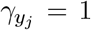, *m < n* results in a higher IDA score because short input durations *D*_*in*_ activate the output more quickly than they inhibit it (Figure 5A). However, increasing the inhibition pathway length *n* is not always advantageous. For example, when *a*_*y*_ = 2, the inhibition pathway amplifies the signal, so longer pathways strengthen inhibition relative to activation regardless of *D*_*in*_. In this regime, the IDA score decreases once *n* exceeds a threshold (e.g., *n* = 4 when *m* = 1). Thus, strong inhibition leads to an optimal inhibition pathway length for a given activation pathway.

**Table 1:**
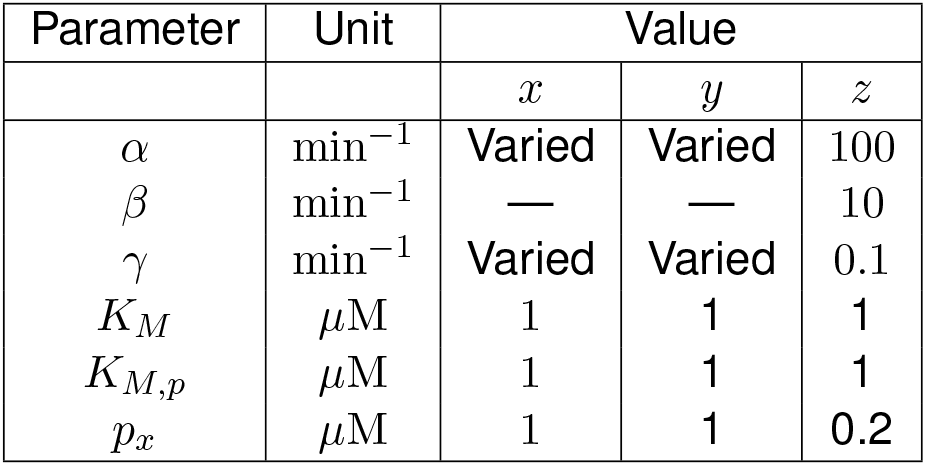
Nominal parameter values for the mathematical models evaluated in Figure 5A to C.

**Table 2:** Nominal parameter values for the mathematical models evaluated in Figure 5D and E.

| Parameter | Unit | Value |  |  |
| --- | --- | --- | --- | --- |
| | | $x$ | $y$ | $z$ |
| $\alpha$ | $\text{min}^{-1}$ | 1 | 1 | Varied |
| $\beta$ | $\text{min}^{-1}$ | — | — | Varied |
| $\gamma$ | $\text{min}^{-1}$ | Varied | Varied | 0.1 |
| $K_M$ | $\mu\text{M}$ | 1 | 1 | 1 |
| $K_{M,p}$ | $\mu\text{M}$ | 1 | 1 | 1 |
| $p_x$ | $\mu\text{M}$ | 1 | 1 | 0.2 |

**Table 3:**
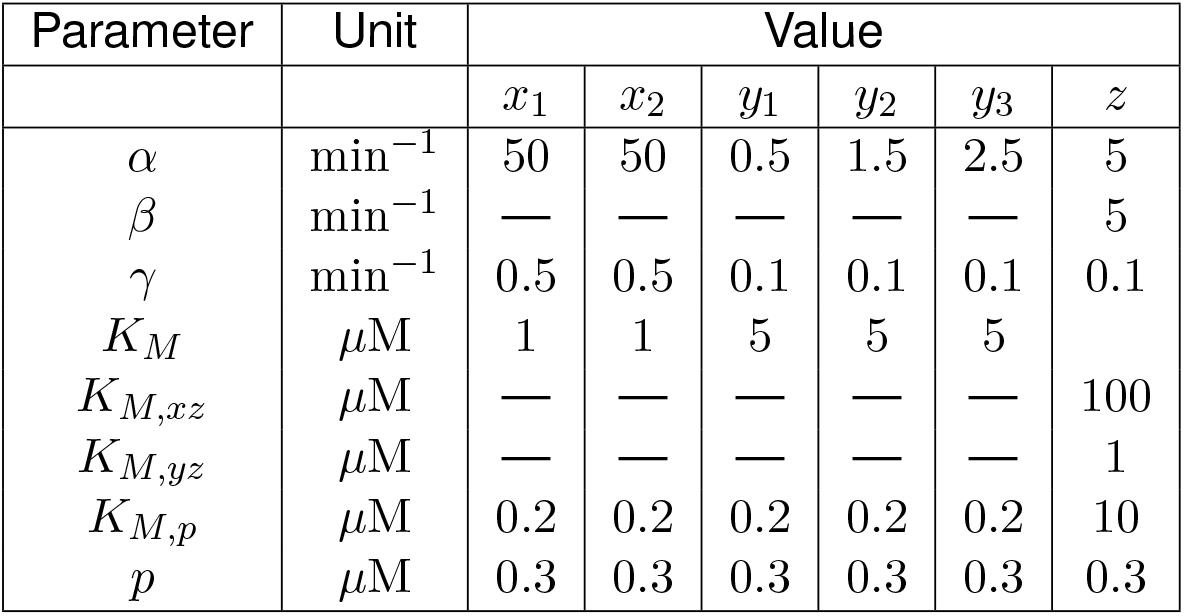
Nominal parameter values for the mathematical models evaluated in Figure 5F and I.

| Parameter | Unit | Value |  |  |  |  |  |
| --- | --- | --- | --- | --- | --- | --- | --- |
| | | $x_1$ | $x_2$ | $y_1$ | $y_2$ | $y_3$ | $z$ |
| $\alpha$ | $\text{min}^{-1}$ | 50 | 50 | 0.5 | 1.5 | 2.5 | 5 |
| $\beta$ | $\text{min}^{-1}$ | — | — | — | — | — | 5 |
| $\gamma$ | $\text{min}^{-1}$ | 0.5 | 0.5 | 0.1 | 0.1 | 0.1 | 0.1 |
| $K_M$ | $\mu\text{M}$ | 1 | 1 | 5 | 5 | 5 | |
| $K_{M,xz}$ | $\mu\text{M}$ | — | — | — | — | — | 100 |
| $K_{M,yz}$ | $\mu\text{M}$ | — | — | — | — | — | 1 |
| $K_{M,p}$ | $\mu\text{M}$ | 0.2 | 0.2 | 0.2 | 0.2 | 0.2 | 10 |
| $p$ | $\mu\text{M}$ | 0.3 | 0.3 | 0.3 | 0.3 | 0.3 | 0.3 |

**Table 4:** Nominal parameter values for the mathematical models evaluated in the Supplemental Figure 9.

| Parameter | Unit | Value |  |  |  |  |  |  |
| --- | --- | --- | --- | --- | --- | --- | --- | --- |
| | | $x_1$ | $x_2$ | $x_3$ | $y_1$ | $y_2$ | $y_3$ | $z$ |
| $\delta$ | $\text{nM hr}^{-1}$ | 20 | 0.2 | 0.3 | 2.5 | 45 | 10 | 0.01 |
| $\epsilon$ | $\text{hr}^{-1}$ | 0.02 | 0.0003 | 0.0003 | 0.0004 | 0.0004 | 0.0003 | 0.001 |
| $K$ | $\text{nM}$ | 0.5 | 60 | 0.6 | 2.2 | 5 | 13 | — |
| $K_{xz}$ | $\text{nM}$ | — | — | — | — | — | — | 8 |
| $K_{yz}$ | $\text{nM}$ | — | — | — | — | — | — | 0.15 |
| $h$ | — | 2 | 3 | 3 | 3 | 4 | 2 | — |
| $h_{xz}$ | — | — | — | — | — | — | — | 2 |
| $h_{yz}$ | — | — | — | — | — | — | — | 4 |

When 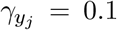 for all *j* ∈ {1, 2, …, *n* }, the decay dynamics of the inhibitory pathway are slower, resulting in more sustained inhibition (Figure 5C). As a result, the IDA score becomes negligible for long pathways (*n* ≥2). Because inhibition is already strong, the cases *a*_*y*_ = 1 and *a*_*y*_ = 2 show little difference. Thus, as in the case *a*_*y*_ = 2 in Figure 5A, excessively strong inhibition makes long inhibitory pathways unfavorable. When *a*_*x*_ = 2, the optimal activation pathway length is *m* = 3 (Figure 5C).

This non-monotonic trend arises from the balance between activation and inhibition strengths (Supplemental Figure 7). For long inputs, increasing *m* mainly extends the output duration with little change in amplitude. For short inputs, increasing *m* initially increases both amplitude and duration, but the response saturates at *m* = 3, after which further increases only prolong the output. Consequently, *m* = 3 yields the highest IDA score.

The previous cases show that comparable activation and inhibition intensities promote higher IDA scores (Figure 5A and 5C). If activation is too strong, both small and large *D*_*in*_ produce long, high-amplitude outputs, lowering the IDA score. If inhibition is too strong, both small and large *D*_*in*_ values produce outputs with small *AUC*, which also reduces the IDA score. When activation and inhibition are more balanced, a more complex pattern emerges, as when 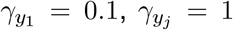 for *j* ∈ {2, …, *n*} (Figure 5B), representing an intermediate case between Figures 5A and 5C.

Figure 5D and E show two circuits with different combinations of *γ*_*j*_ (*j* = 1, …, *n*), each evaluated with four sets of (*α*_*z*_, *β*_*z*_) (additional cases are shown in Supplemental Figure 10). Comparing *α*_*z*_ = 10 (top row) and *α*_*z*_ = 100 (bottom row), the latter produces broader regions with high IDA scores, indicating that a high IDA score requires rapid activation of the output node over a relatively short input duration *D*_*in*_. Among the four *β*_*z*_ values, *β*_*z*_ = 1, 10 yield high IDA scores for cases (E), whereas case (D) favors *β*_*z*_ = 10, 100. Overall, these results highlight the need for a balanced strength between activation and inhibition pathways to obtain the IDA property.

Finally, Figure 5F–I present simulations with a parameter set yielding high IDA and IDD scores (*S*_*IDA*_ = 7.1, *S*_*IDD*_ = 17.8). The parameters, listed in the figure caption, were obtained through random search and are not necessarily optimal. When *D*_*in*_ = 1 min, *X*_2_ activates rapidly while *Y*_3_ remains weak, producing a long-lasting, high-amplitude output. In contrast, when *D*_*in*_ = 20 min, both *X*_2_ and *Y*_3_ activate strongly, generating a pulse with slightly lower amplitude and much shorter duration. A similar response occurs for an even longer input (*D*_*in*_ = 100 min; Supplemental Figure 8). These results suggest that the IFFL acts as a dose filter, selectively responding to transient signals or producing distinct outputs for pulsatile versus sustained inputs^13–17^.

Strictly speaking, the circuit in Figure 5F is a band-pass filter, in the sense that *AUC* as a function of *D*_*in*_ attains a maximum between *D*_*in*_ = 0 and *D*_*in*_ = 1, because the circuit does not produce any output if *D*_*in*_ = 0 (i.e., if there is no input signal). When the input amplitude *u*_0_ is varied instead of *D*_*in*_, *AUC*(*u*_0_) and *D*_*out*_(*u*_0_) behave analogously to *AUC*(*D*_*in*_) and *D*_*out*_(*D*_*in*_) ((the Supplemental Figure 8). Therefore, this circuit behaves as a band-pass filter with respect to both *D*_*in*_ and *u*_0_. This observation is consistent with previous studies reporting that the IFFL circuit can implement a band-pass function and thus detect a certain range of the input dose by producing a larger steady-state response for a certain dose range of a sustained input^14,32,33,49,50^.

## DISCUSSION

We showed that adaptive biomolecular circuits can invert the temporal dose of input signals, producing sustained responses to transient inputs and transient responses to sustained inputs. Analyzing incoherent feedforward loop (IFFL) and negative feedback loop (NFL) motifs, we distinguish two forms of temporal dose inversion: inverse dose duration (IDD) and inverse dose AUC (IDA). While both motifs readily exhibit IDD, IDA arises more frequently in the IFFL due to topology: its activation and inhibition pathways can be tuned independently, enabling low-dose inputs to activate the output without strongly triggering inhibition. In contrast, the NFL couples inhibition to output activation through feedback, limiting such selective responses. Cell-free experiments with a genelet-based IFFL demonstrated IDA behavior. Extending our computational analysis to a broader class of IFFL circuits revealed design rules for temporal dose inversion based on the relative strength of activation and inhibition pathways, and inhibition delay. This insight is consistent with our cell-free experimental findings: changing parameters that modulate output activation and inhibition has a dramatic effect on IDA. Together, these results indicate that temporal dose inversion is a general property of adaptive architectures determined primarily by network topology, while also depending on underlying biochemical implementations and parameter values. Our experimental demonstration uses a genelet-based IFFL^21^, but other programmable *in vitro* platforms, such as the polymerase-exonuclease-nickase (PEN) toolbox^51^ and cell-free gene networks^52^, could be adopted for the same purpose. Such components could expand synthetic signaling systems capable of processing and decoding pulsatile inputs^50,53,54^.

Our finding that the IFFL motif acts as a dose filter aligns with several studies suggesting its capacity to detect pulsatile signals^13–17^. However, prior work did not systematically examine how dose–response inversion arises or how it enables pulse detection. Moreover, pulse detection principles outlined in previous work differ from those we identified here. For example, Lormeau et al.^15^ detected single pulses using an interlocked IFFL, and Benzinger et al.^14^ detected pulse trains with a diamond-shaped IFFL. Both rely on falling-edge detection: while the input is present, the output remains minimal because activation and inhibition are triggered simultaneously; once the input disappears, the output increases as the activation signal decays more slowly. In contrast, in our mechanism the output rises immediately after input onset and decays once the inhibition pathway dominates. Because triggering the inhibition pathway requires a sufficient input dose, shorter or weaker inputs stimulate the output more effectively, producing dose filtering through temporal dose inversion. A related strategy was proposed for pulse-train detection in^17^, although that model relied on an extremely high Hill coefficient, limiting its biological realism.

The ability of IFFL motifs to generate temporal dose inversion relates to their well-known band-pass filtering properties^32,33,50,55^, where independently tuned activation and inhibition thresholds restrict responses to a defined input range. In contrast, temporal dose inversion evaluates dose through time-dependent input features. Our simulations reveal a “duality” between input duration and amplitude (Figure 2D): temporal dose inversion emerges both when duration varies at constant amplitude, and when amplitude varies at constant duration. Accordingly, dose filtering can be interpreted as duration filtering or amplitude filtering when one input dimension is held constant. This duality may underlie the broad utility of IFFL motifs, including circuits that detect regions where molecule concentrations fall within a specific range leading to spatial patterning in bacteria^32^ and mammalian cells^33^.

The concept of temporal dose inversion may help describe and interpret behaviors associated with temporal coding in natural living systems^4,56^. For example, tumor necrosis factor (TNF)-induced NF-*κ*B signaling exhibits responses that depend on the temporal properties of TNF pulses. First, TNF stimulates both the pro-apoptotic caspase pathway and the pro-survival NF-*κ*B-driven gene transcription pathway, which in turn inhibits some steps in the pro-apoptotic caspase signaling, leading to a topology consistent with the IFFL motif. Lee et al.^57^ reported that a 1-min TNF pulse induces apoptosis more effectively than a 1-hour TNF pulse. This behavior may be related to temporal dose inversion. More broadly, cells interpret multiple temporal features of inputs, such as amplitude, duration, and frequency^3^. Exploring temporal dose inversion across various input features may provide insights into how biological systems decode temporally varying signals.

Temporal dose inversion circuits could expand and simplify the combinatorial recognition of multiple temporal features of a signal, similar to strategies observed in living systems. For example, Benzinger et al.^14^ demonstrated pulse-train signal processing based on the amplitude and duty cycle using two readouts. Temporal dose-response may simplify the design of circuits that detect duty cycle or perform complex temporal signal processing. Libraries of circuits for temporal coding and detection may expand our knowledge of molecular communication, with potential relevance for diagnostics^58^, therapeutics^59^, and material development^60^.

## Supplemental information

### Linear systems do not exhibit temporal dose inversion

We consider causal linear time invariant systems with input *u*(*t*) and state/output *y*(*t*), characterized by a non-negative impulse response function *h*(*t*). This class of systems is relevant to our study, because any physically plausible biological model must present non-negative variables at all times.

Assume *y*(0) = 0 and *u*(*t*) = 0 for *t <* 0. The forced response of this type of system is:

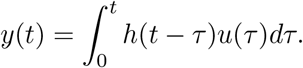

Consider a finite-duration step input *u*_*T*_ (*t*):

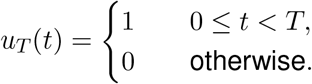

The response to the step input with *u*(*t*) = 0 for *t* < 0 and *u*(*t*) = 1 for *t* ≥ 0 is

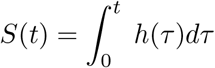

and it is non-decreasing, because *h* is non-negative.

Input *u*_*T*_ can be seen as the difference between the unit step at zero (*u*(*t*) = 0 for *t* < 0 and *u*(*t*) = 1 for *t*≥ 0) and the unit step at *T* (*u*(*t*) = 0 for *t* < *T* and *u*(*t*) = 1 for *t*≥ *T*). In view of linearity, the response to the difference of the two signals is difference of the responses to the two signals:

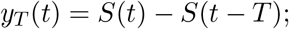

note that, for *t < T*, we have *y*_*T*_ (*t*) = *S*(*t*).

Now, consider two inputs 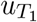 and 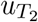, with *T*_1_ < *T*_2_. Since *S* is non-decreasing, we have that *S*(*t* − *T*_1_) ≥ *S*(*t* − *T*_2_). Hence

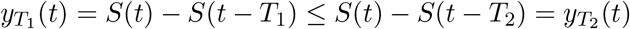

This means that the magnitude of the response *y*(*t*) scales with the duration of the input signal. In other words, this kind of system does not admit temporal dose inversion.

### Sensitivity of temporal responses to model parameters in the simple IFFL and NFL models

**Figure 6:**
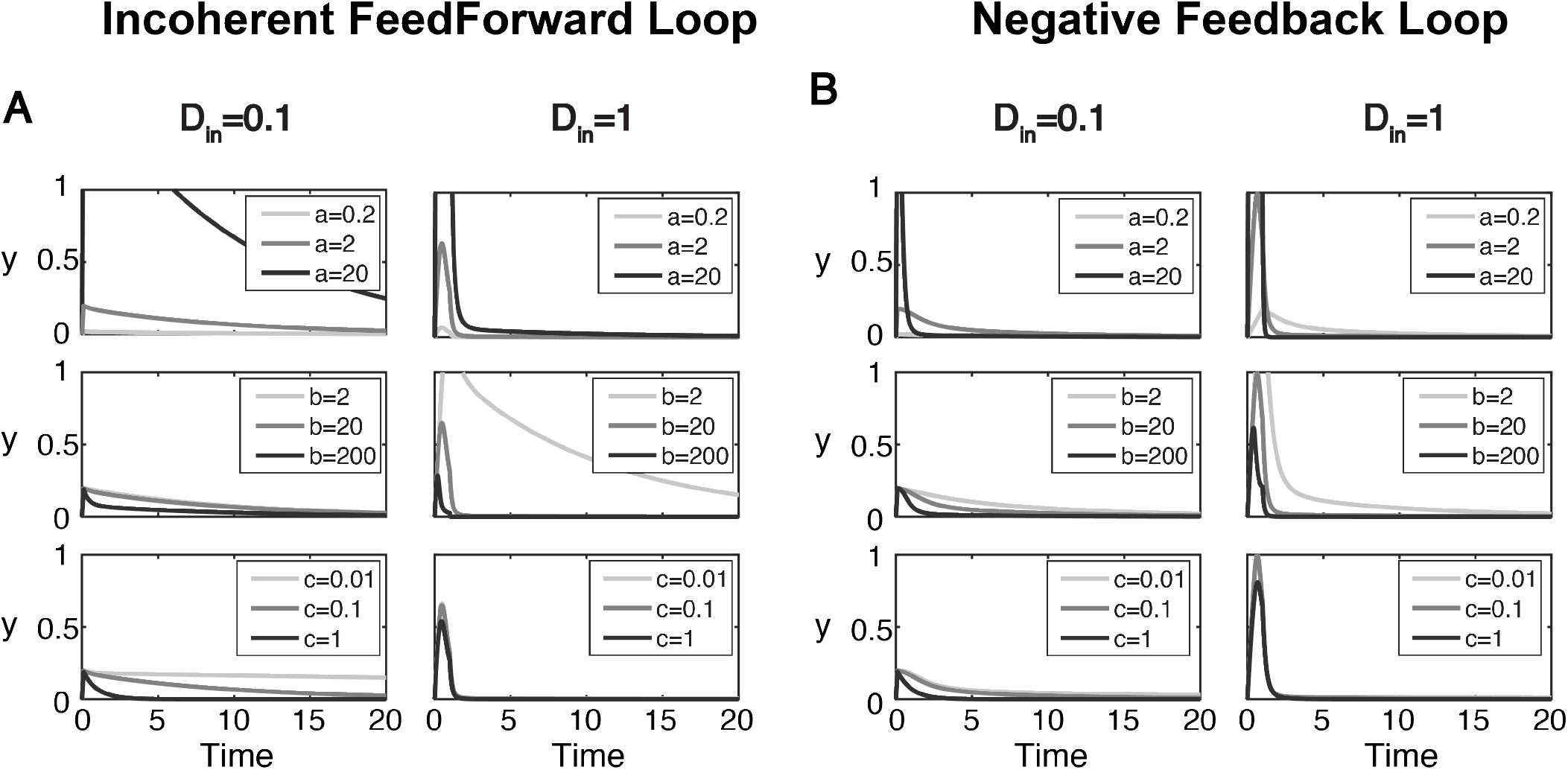
Evaluating the influence of individual parameter variations on the temporal response of the IFFL and NFL circuits. Temporal responses of (A) the IFFL and of (B) the NFL circuit to a rectangular input with amplitude *u*_0_ = 1 when *D*_*in*_ = 0.1 (left) and when *D*_*in*_ = 1 (right). The values of varying parameters are indicated in the legends. All other parameters are kept constant at their nominal value *a* = 2, *b* = 20, *c* = 0.1, *p* = 2.

### Circuit dynamics with different combinations of (*m, n*)

**Figure 7:**
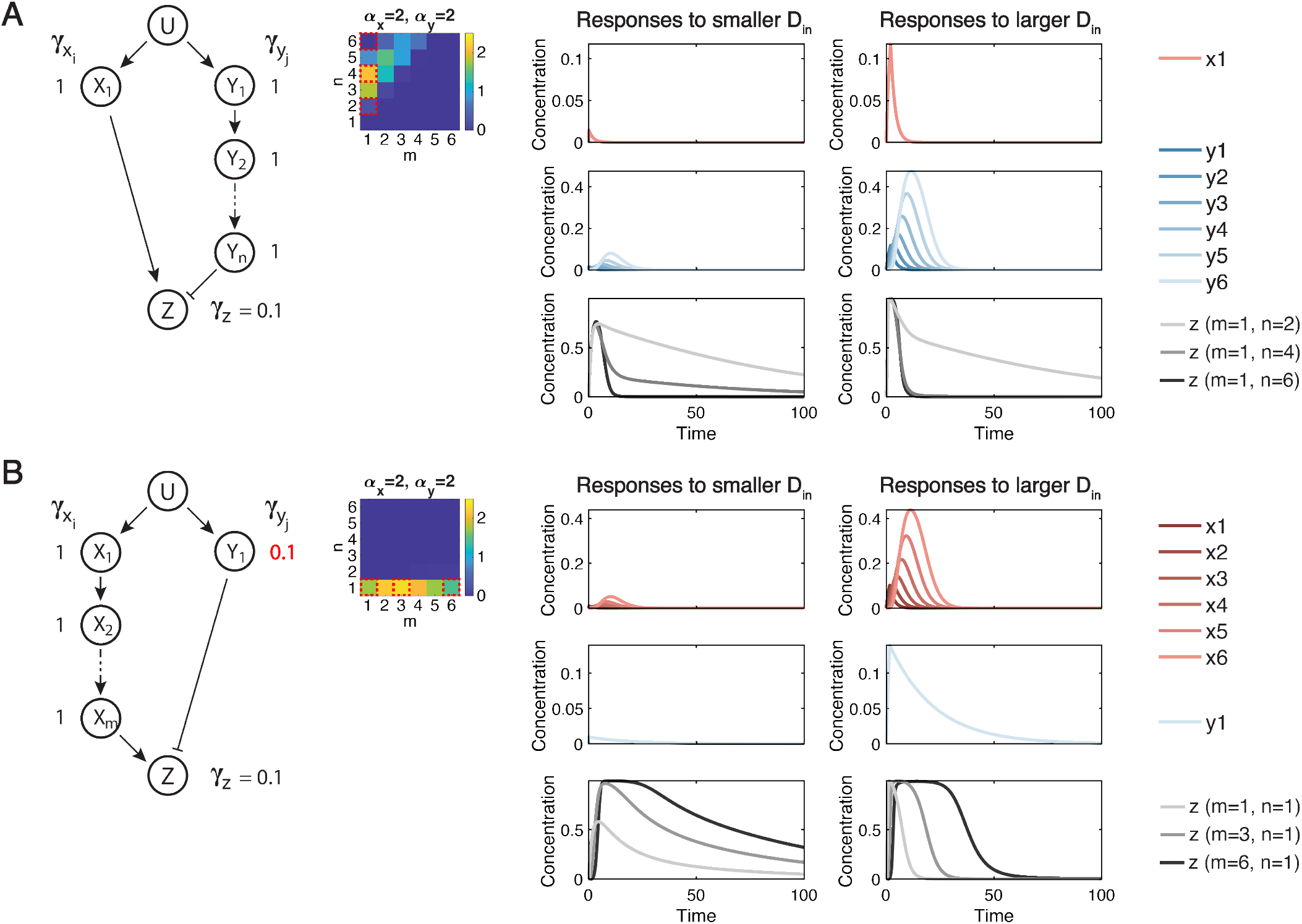
Dynamics of the circuit associated with the model in Equations 13-17 with different combinations of (*m, n*). (A) Behaviour of the circuit in Figure 5A, having 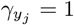 for all *j*, with *m* = 1 and different values of *n*. (B) Behaviour of the circuit in Figure 5C, having 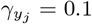 for all *j*, with *n* = 1 and different values of *m*.

### Additional details of the signaling IFFL circuit showing a high IDA score

**Figure 8:**
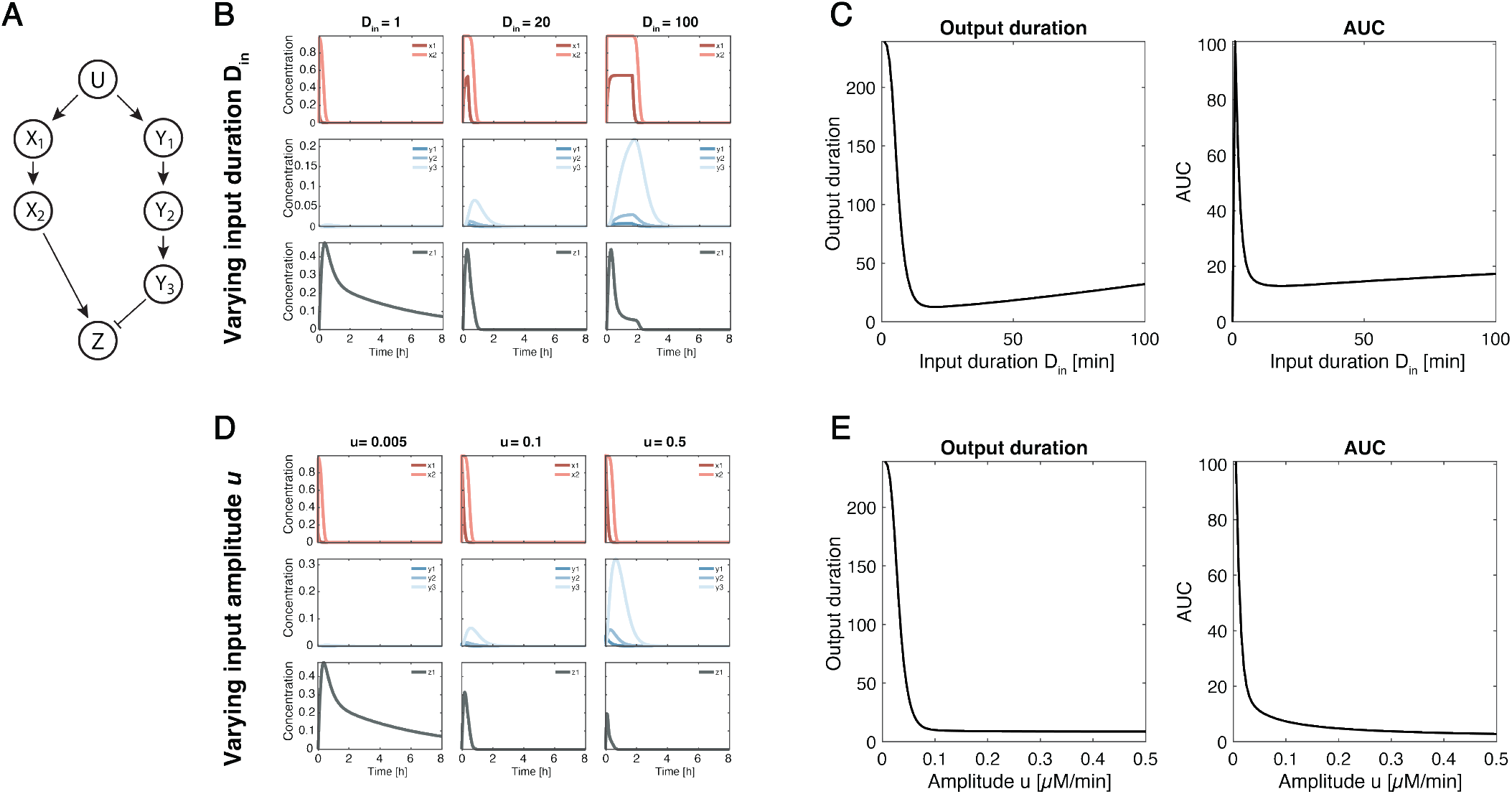
Details of the signaling IFFL circuit shown in Figure 5F to I. (A) Diagram of the signaling IFFL circuit. The circuit has the parameters listed in Table 3. (B) Dynamics of each node with *D*_*in*_ = 1 min (left), *D*_*in*_ = 20 min (center) and *D*_*in*_ = 100 min (right). (C) Output duration and *AUC* as a function of *D*_*in*_ with *u*_0_ = 0.005. (D) Dynamics of each node with *u* = 0.005 µMmin^*−*1^ (left), *u* = 0.1 µMmin^*−*1^ min (center) and *u* = 0.5 µMmin^*−*1^ (right). (E) Output duration and *AUC* as a function of *u* with *D* = 1 min

### A GRN circuit showing the IDA property

**Figure 9:**
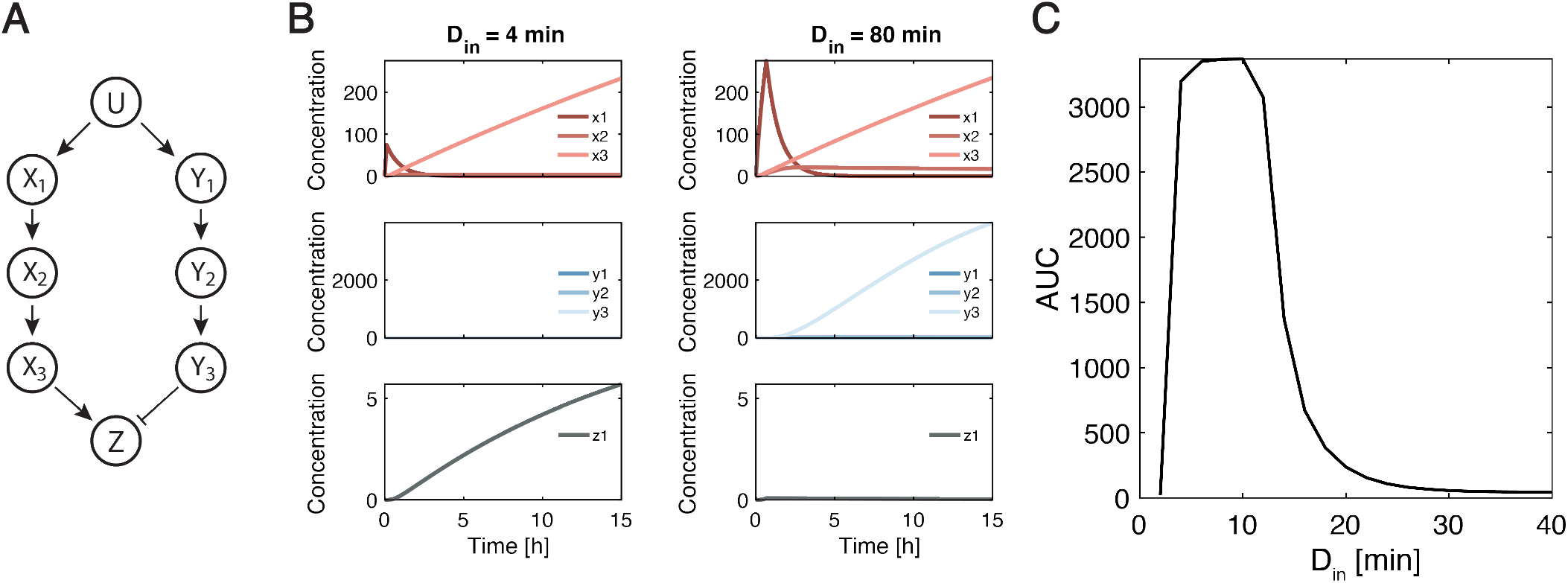
A GRN IFFL circuit showing the IDA property; the mathematical model is described in STAR Methods 4. (A) Diagram of the GRN IFFL circuit. (B) Dynamics of each node with *D*_*in*_ = 4 min (left) and *D*_*in*_ = 80 min (right). The input amplitude was kept at *u*_0_ = 0.5 (C) *AUC* as a function of *D*_*in*_. Parameter values are listed in Table 4.

### Activation and inhibition rates on the output node affects IDA property

In the main text, we evaluated how the parameters of node *X*_*i*_ (*i* = 1, …, *m*) and *Y*_*j*_ (*i* = 1, …, *n*) affect the IDA property of generalized IFFL circuits. Here, we see how the parameters of the output node *Z* affect the IDA property of the same circuit motifs. Figure 10 shows three cases (different combinations of *γ*_*j*_ (*j* = 1, …, *n*)) as evaluated in Section 4. For each case, eight different sets of (*α*_*z*_, *β*_*z*_) are evaluated. When we compare *α*_*z*_ = 10 (top row) and *α*_*z*_ = 100 (bottom row) for each case, *α*_*z*_ = 100 gives broader areas with high IDA score for all the cases. These results suggest that, for obtaining a high IDA score, the output node has to be quickly activated during a relatively shorter period with a smaller *D*_*in*_. Among four different values of *β*_*z*_, *β*_*z*_ = 1, 10 give high IDA scores for case (B) and (C), while case (A) prefers *β*_*z*_ = 10, 100. These results supports the argument that relative balance of the activation and inhibition determines the IDA property, which was stated in the main text.

**Figure 10:**
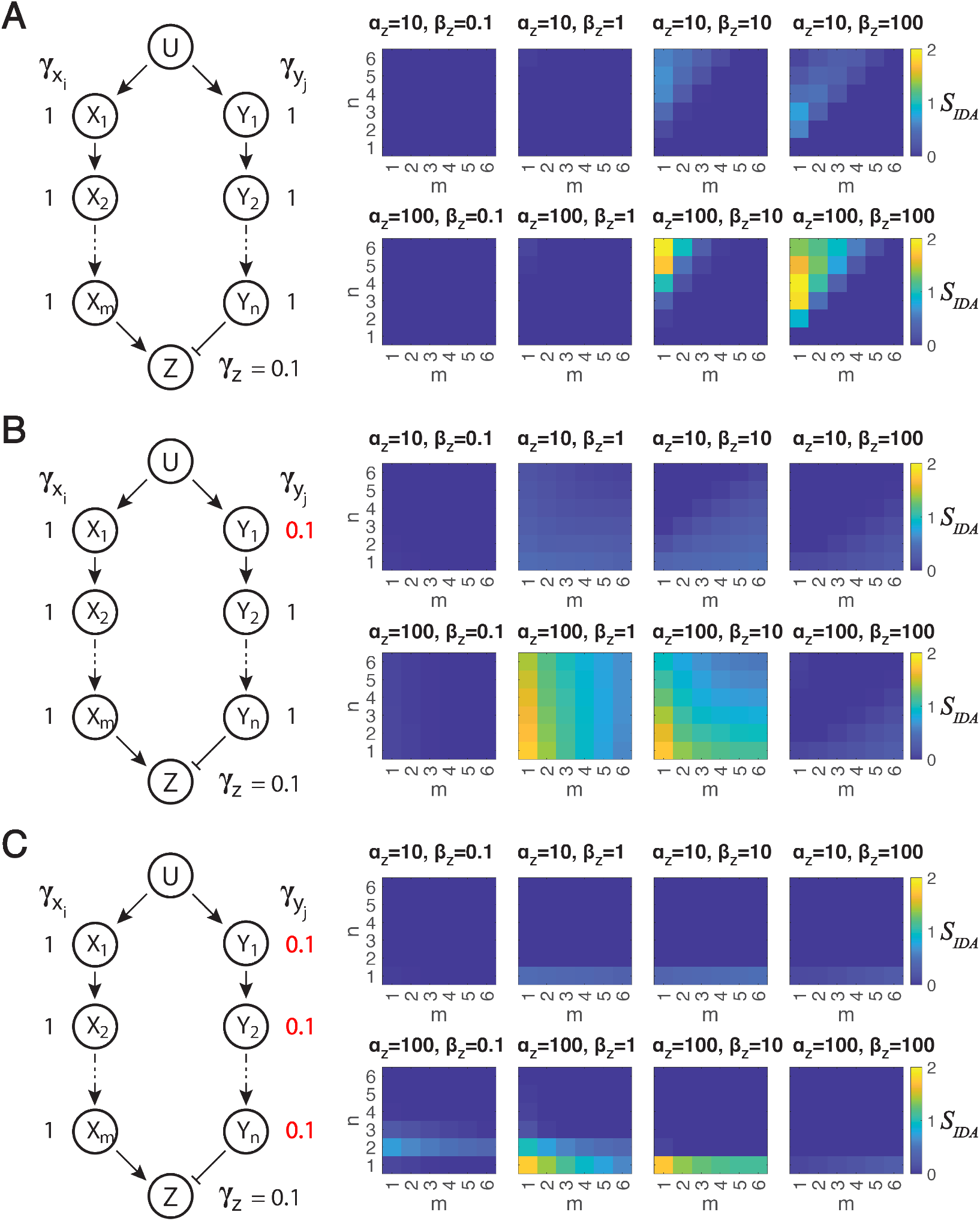
Parameters of the output node *Z* affects the IDA property of the generalized IFFL circuits. (A to C) Each panel shows the circuit diagram of a generalized IFFL network, along with heat maps of IDA scores *S*_*IDA*_ in the *m*-*n* plane for different values of *α*_*z*_ and *β*_*z*_, reported above each heat map. The basal deactivation rates are reported next to each node in the circuit diagram: in all three cases, 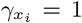 for all *i* = 1, …, *m*, and moreover (A) 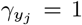 for all *j* = 1, …, *n*; (B) 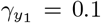, and 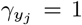 for *j* = 2, …, *n*; (C) 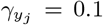 for all *j* = 1, …, *n*. When not varied, the parameters are kept constant at their nominal values: *α*_*x*_ = 1, *α*_*y*_ = 1, *γ*_*z*_ = 0.1, *K*_*M,x*_ = *K*_*M,y*_ = *K*_*M,z*_ = 1.0, *p*_*x*_ = *p*_*y*_ = 1.0, *p*_*z*_ = 0.2, *u*_0_ = 0.1; *D*_*in*_ is logarithmically varied from *D*_*in*_ = 0.1 to *D*_*in*_ = 10.

**Figure 11:**
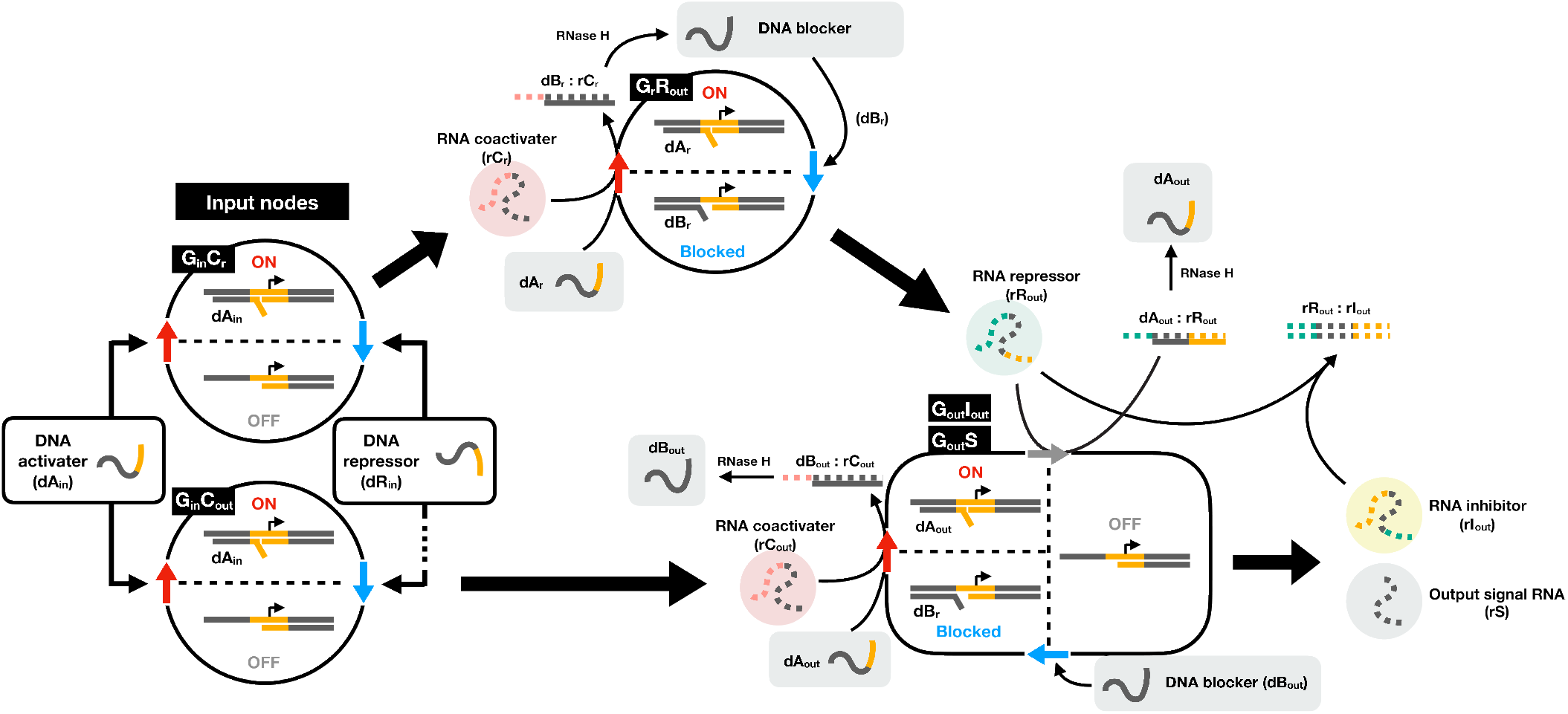
A detailed schematic of the genelet-based IFFL circuit. Each genelet node is depicted either as a circle (input and intermediate nodes: *G*_*in*_*C*_*out*_ or *G*_*in*_*C*_*r*_, *G*_*r*_ *R*_*out*_) or as a rounded rectangle (output node: *G*_*out*_*S*). Genelets are switched ON (red arrows) or OFF (blue arrows) mediated by RNA coactivators or repressors respectively, which are indicated by colored circles. Single-stranded DNA species are indicated by gray rounded rectangles.

**Figure 12:**
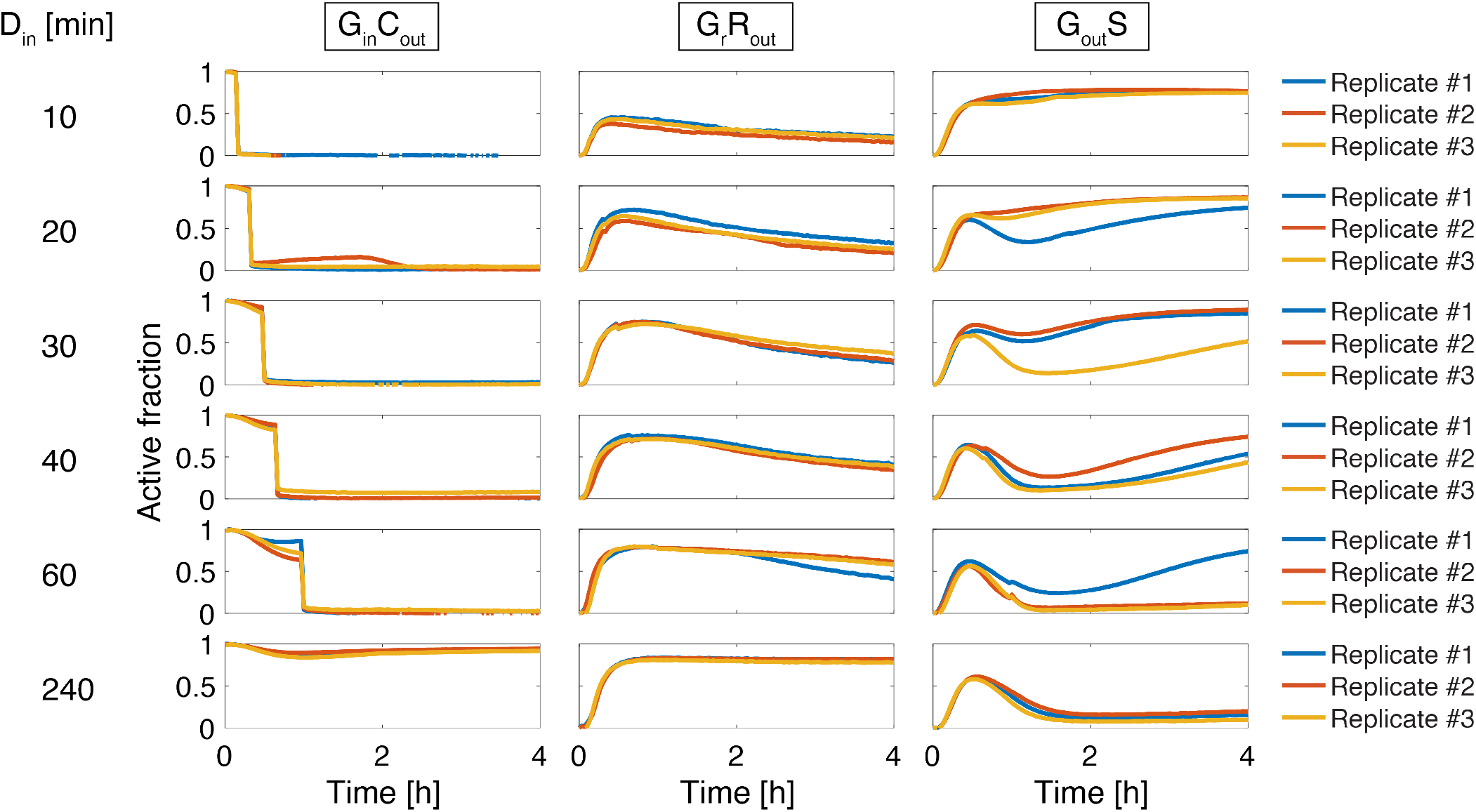
All the replicates of the genelet IFFL experiments with varying input durations. We have separate experiments for six different input durations: 10, 20, 30, 40, 60, 240 min. Experiments for each input duration have three replicates, and the active fractions of G_in_C_out_, G_r_R_out_, and G_out_S are shown for each replicate.

**Figure 13:**
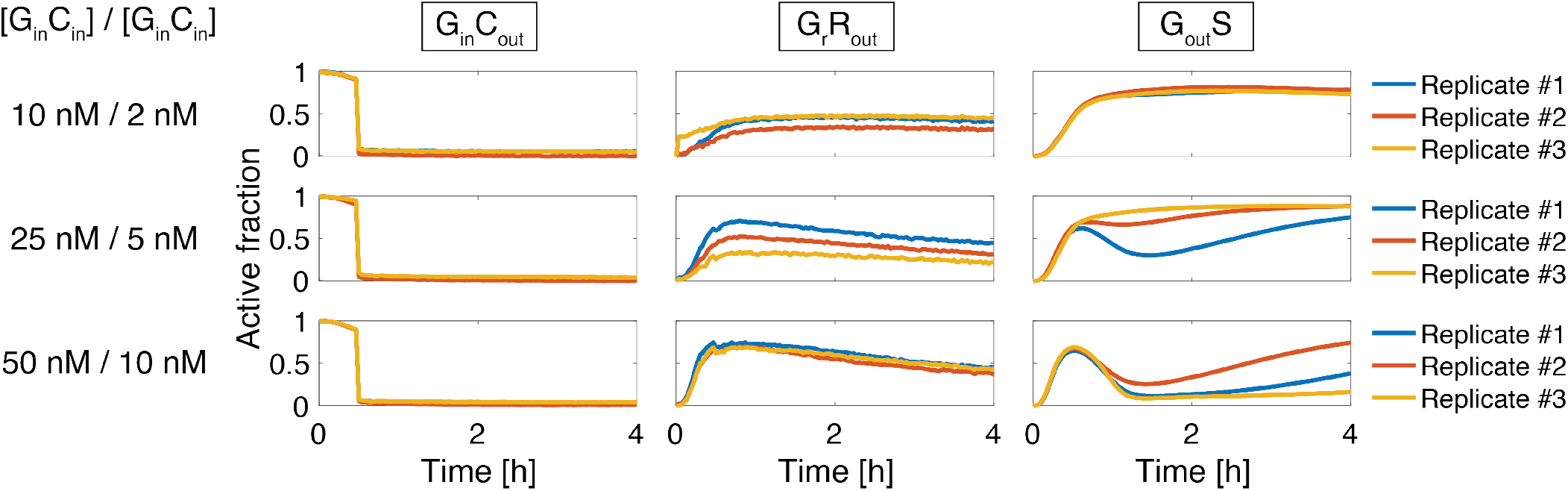
All the replicates of the genelet IFFL experiments with varying input genelet concentrations. We have separate experiments for three different sets of [G_in_C_out_]/[G_in_C_r_]: 10 nM / 2 nM, 25 nM / 5 nM, 50 nM / 10 nM. Experiments for each input genelet concentrations have three replicates, and the active fractions of G_in_C_out_, G_r_R_out_, and G_out_S are shown for each replicate.

### Detailed architecture of the genelet-based IFFL circuit

Our genelet-based IFFL is derived from the design originally developed by Schaffter and coworkers^21^. We modified their design by incorporating an additional genelet, *G*_*out*_*I*_*out*_, at a low conentration (1 nM) to suppress the effect of transcription leakage from *G*_*r*_*R*_*out*_, adjusting component concentrations to reinforce the temporal dose inversion, and employing a different set of input genelet nodes to enable sharper ON/OFF switching. In the original genelet notation of Schaffter et al., the genelets *G*_*in*_*C*_*out*_, *G*_*in*_*C*_*r*_, *G*_*r*_*R*_*out*_, *G*_*out*_*I*_*out*_, *G*_*out*_*S* in our implementation correspond to *G*8*C*1, *G*8*C*3, *G*3*R*1, *G*1*I*1, *G*1*S*, respectively. A detailed schematic of the resulting circuit architecture is shown in the Supplemental Figure 11.

### Complete experimental dataset of genelet IFFL experiments

All the replicates for the genelet IFFL experiments with varying input durations are shown in the Supplemental Figure 12. We have conducted separate experiments to measure responses of the genelet-based IFFL circuit to six different input durations: 10, 20, 30, 40, 60, 240 min. For each input duration, we obtained three replicates to evaluate the variance of the experiment.

All the replicates for the genelet IFFL experiments with varying input genelet concentrations are shown in the Supplemental Figure 13. We have conducted separate experiments for three different sets of [G_in_C_out_] / [G_in_C_r_]: 10 nM / 2 nM, 25 nM / 5 nM, 50 nM / 10 nM. The input duration was fixed at 30 min. For each input concentration, we obtained three replicates to evaluate the variance of the experiment.

For both experiments, the variation of G_out_S active level is correlated with the variation of G_r_R_out_ active level; the higher active fraction of G_r_R_out_ is, the lower the active fraction of G_out_S. The variation of G_r_R_out_ is most likely due to the fluctuation of enzyme concentrations and activities.

### A mathematical model of the genelet IFFL

Here we describe the detailed mathematical model of the genelet IFFL that we designed for the biological realization of the IDA property in Section 2. We use the following mathematical model for the genelet IFFL, which is exactly the same as the model established by Schaffter et al.^21^:

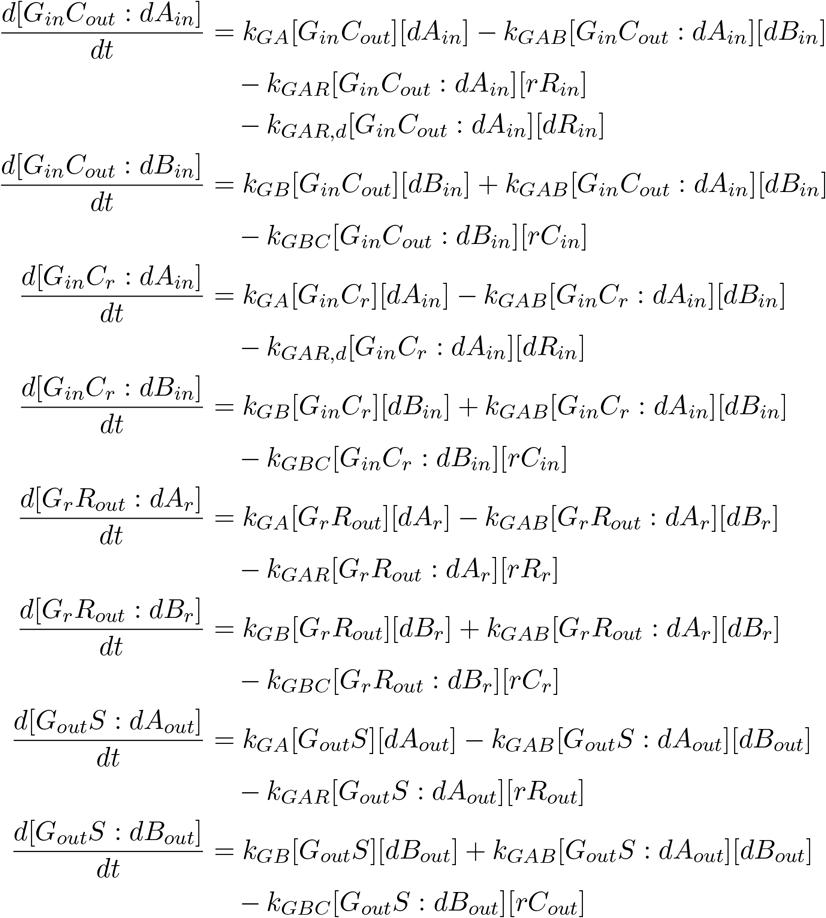

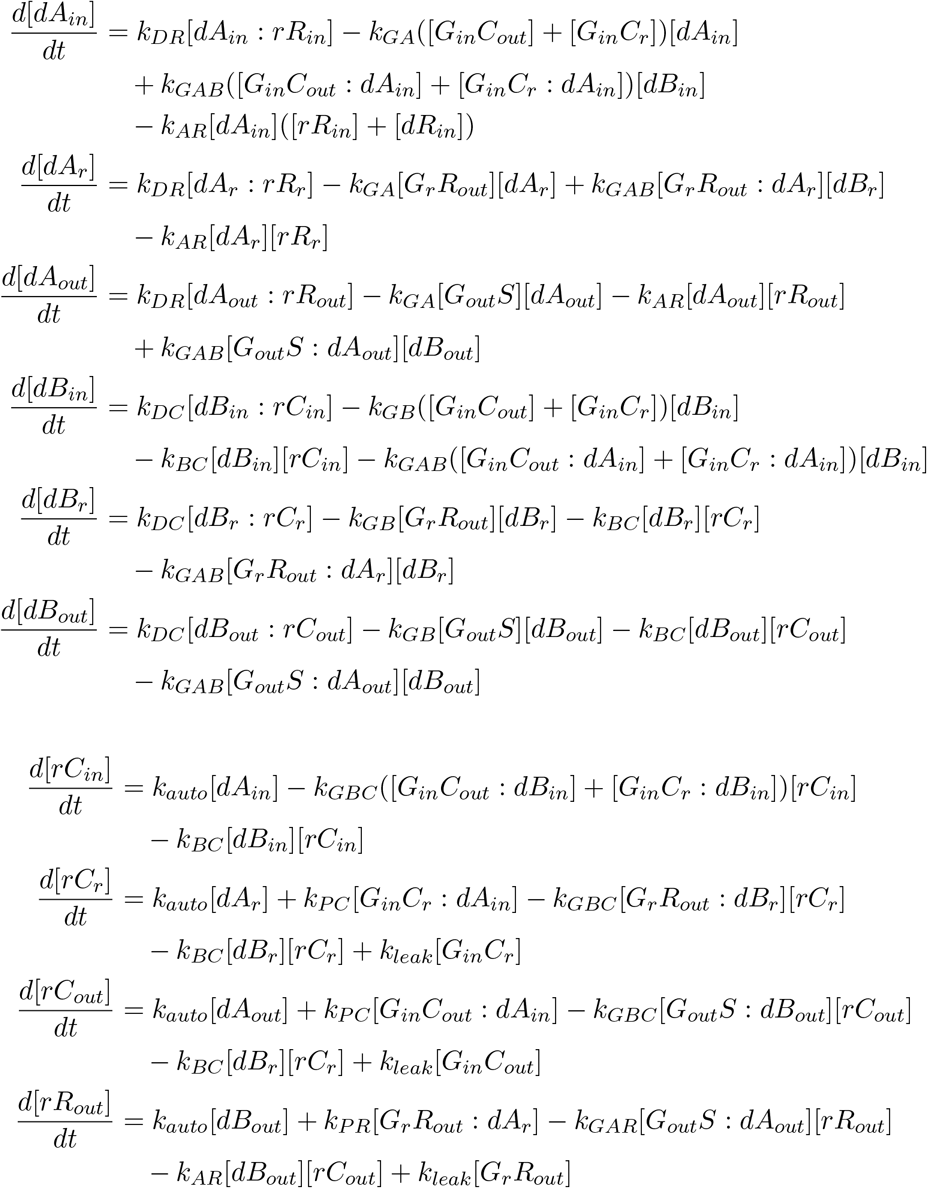

where *k*_*GA*_, *k*_*GAB*_, *k*_*GAR*_, *k*_*GBC*_, *k*_*AR*_, *k*_*BC*_ are the binding constants between genelets and DNA activators, activated genelets and RNA repressors, blocked genelets and RNA coactivators, DNA activators and RNA repressors, DNA blockers and RNA coactivators respectively; *k*_*GAR,d*_ is the binding constant between activated genelets and DNA repressor; *k*_*P C*_, *k*_*P R*_ are the production rates of RNA coactivators and repressors respectively; *k*_*DC*_, *k*_*DR*_ are the degradation rates of DNA-RNA complexes. Here we assume binding constants and rate constants are identical for all the genelet nodes; *k*_*auto*_ is the production rate of RNA repressor or coactivator from their corresponding DNA activator or DNA blocker. The produced RNAs cause auto-inhibition or auto-activation, which has been reported by Schaffter et al.^21^; *k*_*leak*_ is the production rate of RNA repressor or RNA coactivator from their corresponding genelets at OFF state or BLOCKED state.

We also have the following mass balance equations:

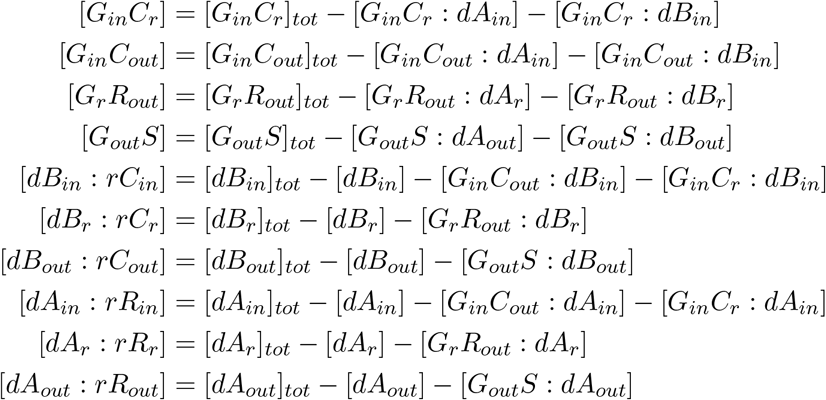

The parameter values fitted based on the experimental results are listed in Table 9,10, 11 and 12.

### Sample-to-sample variability induced by enzymatic parameter fluctuations

We hypothesize that the experimental fluctuations observed in Figure 3 is primarily originated from pipetting variability specifically for enzyme solutions, as the enzyme solution contains a high concentration of glycerol, which alters its viscosity and wetting properties compared to aqueous buffers, making it more prone to pipetting variability. To support this hypothesis, we performed a Monte Carlo analysis in which enzymatic parameters were stochastically varied.

Specifically, we considered variability in two groups of kinetic parameters: (i) transcription-related parameters *k*_*pc*_, *k*_*pr*_, *k*_*pi*_ and (ii) RNA degradation-related parameters *k*_*dc*_, *k*_*dr*_. For each group, parameters are scaled by its common perturbation factor, which is associated with the pipetting fluctuation and batch-to-batch activity variation of T7 RNA polymerase and RNase H, which are independent of genelet nodes. We defined the perturbation factor as follows:

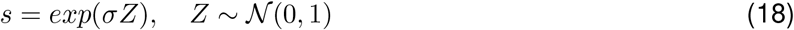

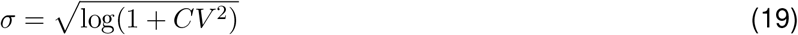

where *σ* is the standard deviation of the underlying normal distribution in log space, and *CV* is the coefficient of variation of the resulting perturbation factor *s. CV* was set to 0.1. All remaining parameters are fixed at the nominal parameter set (Table 6), and at the fitted circuit parameters (Table 9, 10, and 11). Input pulse durations of 10, 20, 30, 40, 60, 240 min were tested. For each Monte Carlo sample, the ODE model of the genelet-based IFFL was simulated for four hours, and the area under the curve (*AUC*) of *G*_*out*_*S* was computed from the obtained time evolution. For each input duration, 500 independent simulations were performed. The obtained results are shown in Figure 14. The simulated standard deviations are compared with those observed experimentally, supporting the hypothesis that the experimental fluctuation can be explained by the fluctuations of the enzymatic activity derived from pipetting and preparation variability.

**Figure 14:**
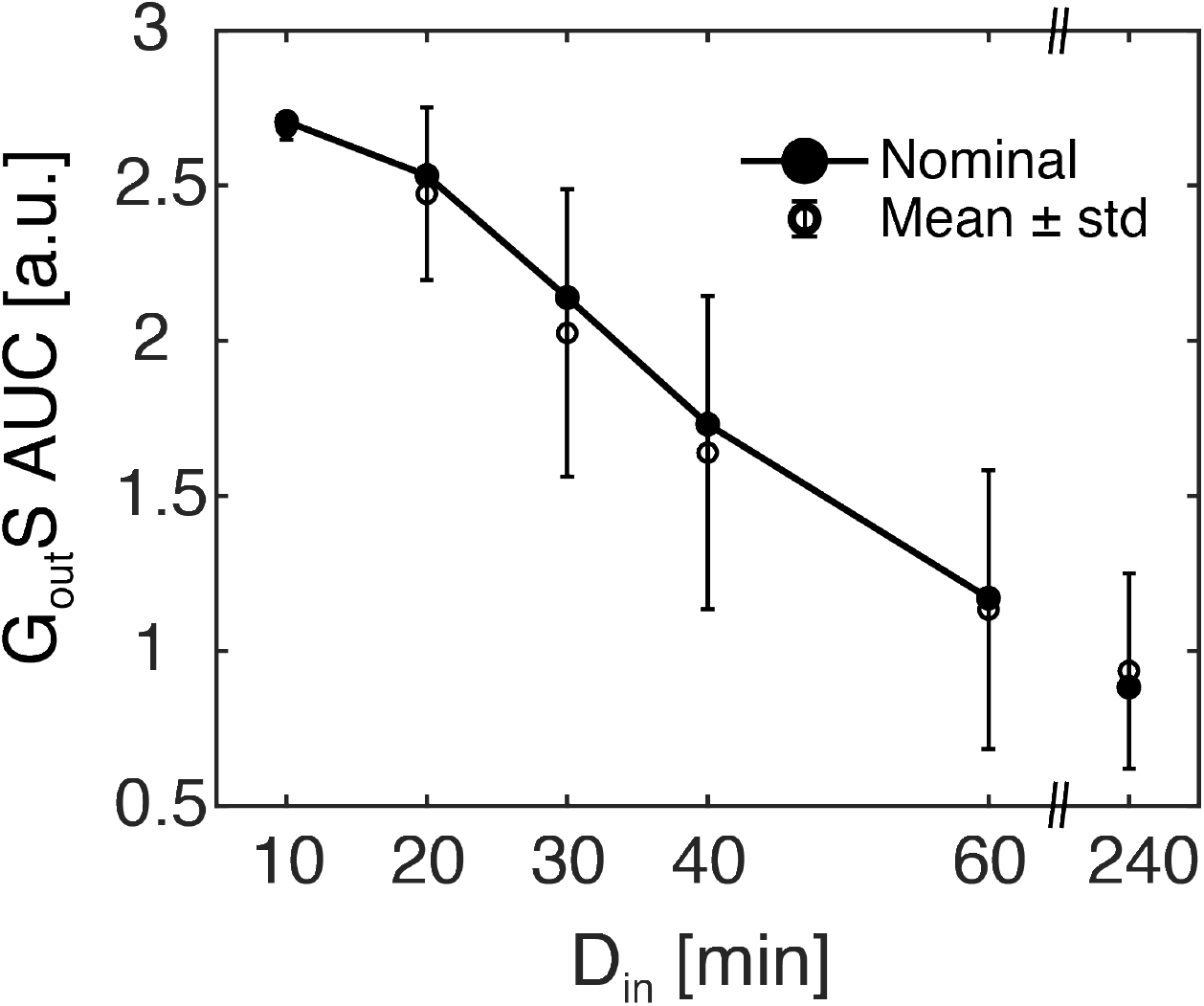
Monte Carlo analysis quantifies the sample-to-sample variability of the output *AUC* of the genelet-based IFFL induced by the fluctuations in enzymatic parameters.

### Analysis of additional adaptive circuits

In the main text, our focus was on the IFFL and NFL. However, there are other types of adaptive circuit motifs. Here, we briefly evaluate two additional circuit motifs: IFFL combined with NFL (hereinafter, referred to as IFFL+NFL), and an antithetic integral feedback (Figure 15). Circuit motifs in real biological networks are often intertwined, which suggests evaluating combined circuit motifs^20,61^. Here, we only evaluate IFFL+NFL as a combined motif because it is the only circuit pattern of two circuit nodes (not including the input node) that has both IFFL and NFL motif. The ODE model of IFFL+NFL is defined as

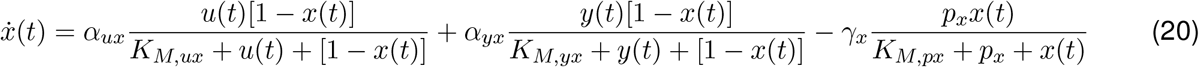

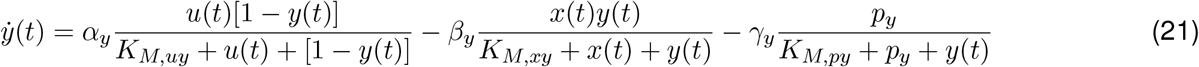

**Figure 15:**
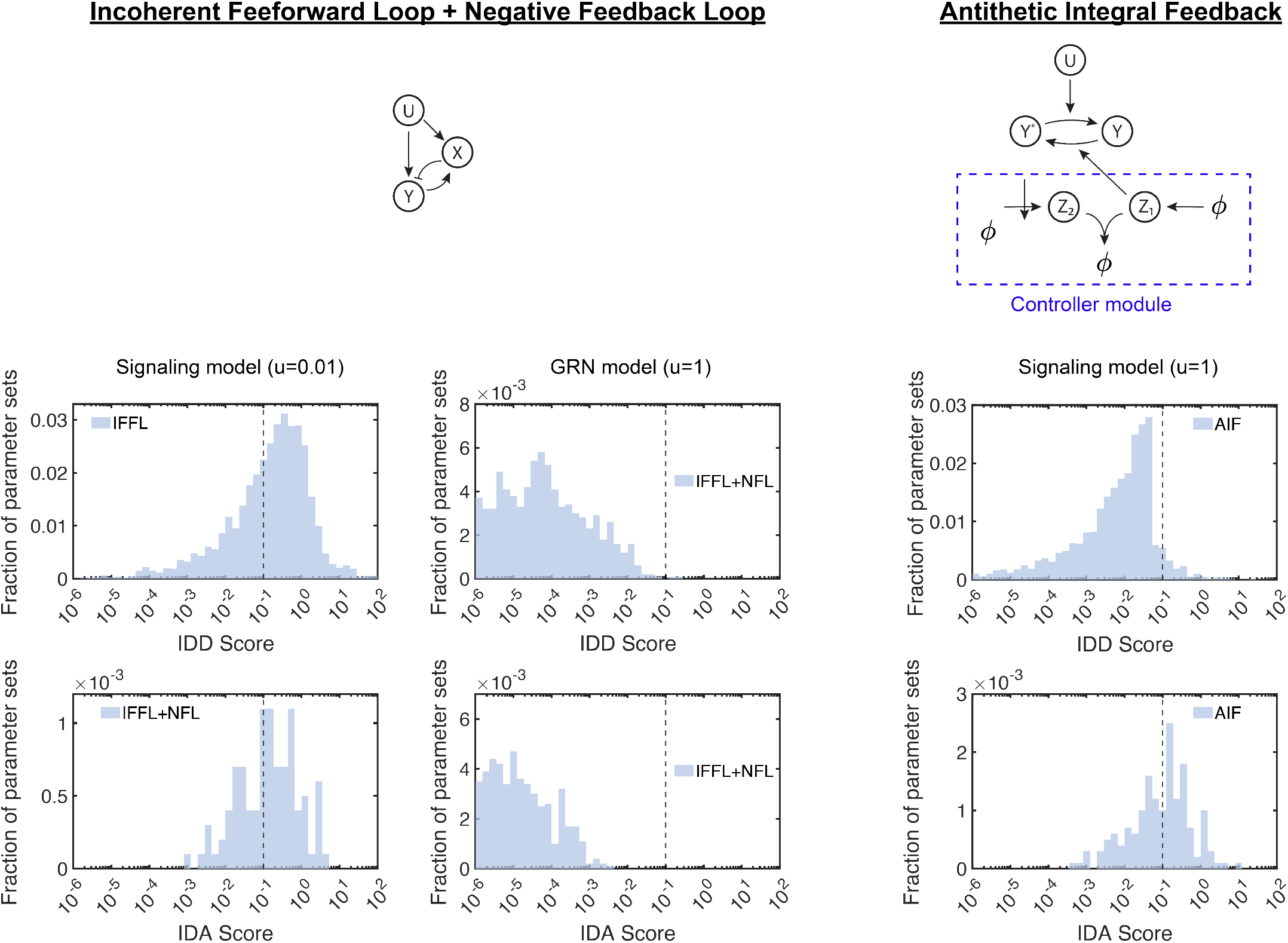
Histograms showing the occurrence of positive IDD and IDA scores in the IFFL+NFL circuit and the antithetic integral feedback circuit. The selected input amplitudes are *u* = 0.01 for the signaling IFFL+NFL, *u* = 1 for the GRN IFFL+NFL, and *u* = 0.01 for the antithetic integral feedback.

In the parameter set evaluated here, IFFL+NFL showed an intermediate histogram between IFFL and NFL. We also evaluated the effect of an antithetic integral feedback. The antithetic integral feedback is a powerful controller that realizes a robust perfect adaptation. Even though it is still unclear how adaptation and the dose inversion properties are related, it is an intriguing question how the antithetic integral feedback affects the dose inversion properties. In the antithetic integral feedback controller, we introduce two additional species, *z*_1_ and *z*_2_, to regulate the system of interest. *z*_1_ is constitutively produced at a constant rate *µ*, whereas the production of *z*_2_ is activated by the inactive form of *y* with rate constant *θ*. Species *z*_1_ and *z*_2_ bind to each other to form an inactive complex. Both *z*_1_ and *z*_2_ are also subject to basal degradation. In this study, the antithetic integral feedback controller was applied only to the signaling model, because the binding reaction between *z*_1_ and *z*_2_ can be naturally interpreted as a protein-protein interaction. Although it is possible to combine gene regulation with the antithetic integral feedback controller, exploring such implementation is beyond the scope of this work. The ODE model of the antithetic integral feedback circuit is as follows:

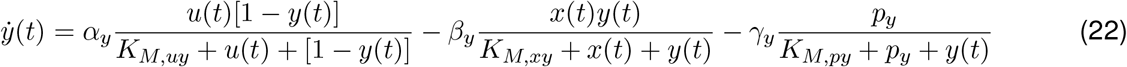

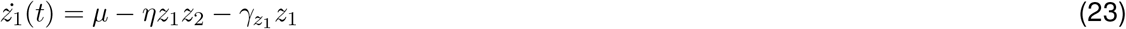

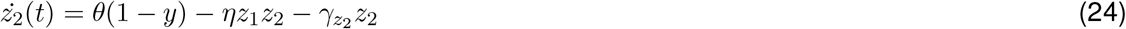

### The occurrence of temporal dose inversion properties depends on input amplitude

Figure 16 shows a complete set of histograms of IDD and IDA scores obtained by the random parameter sampling for the signaling and the GRN models of the IFFL and NFL.

**Figure 16:**
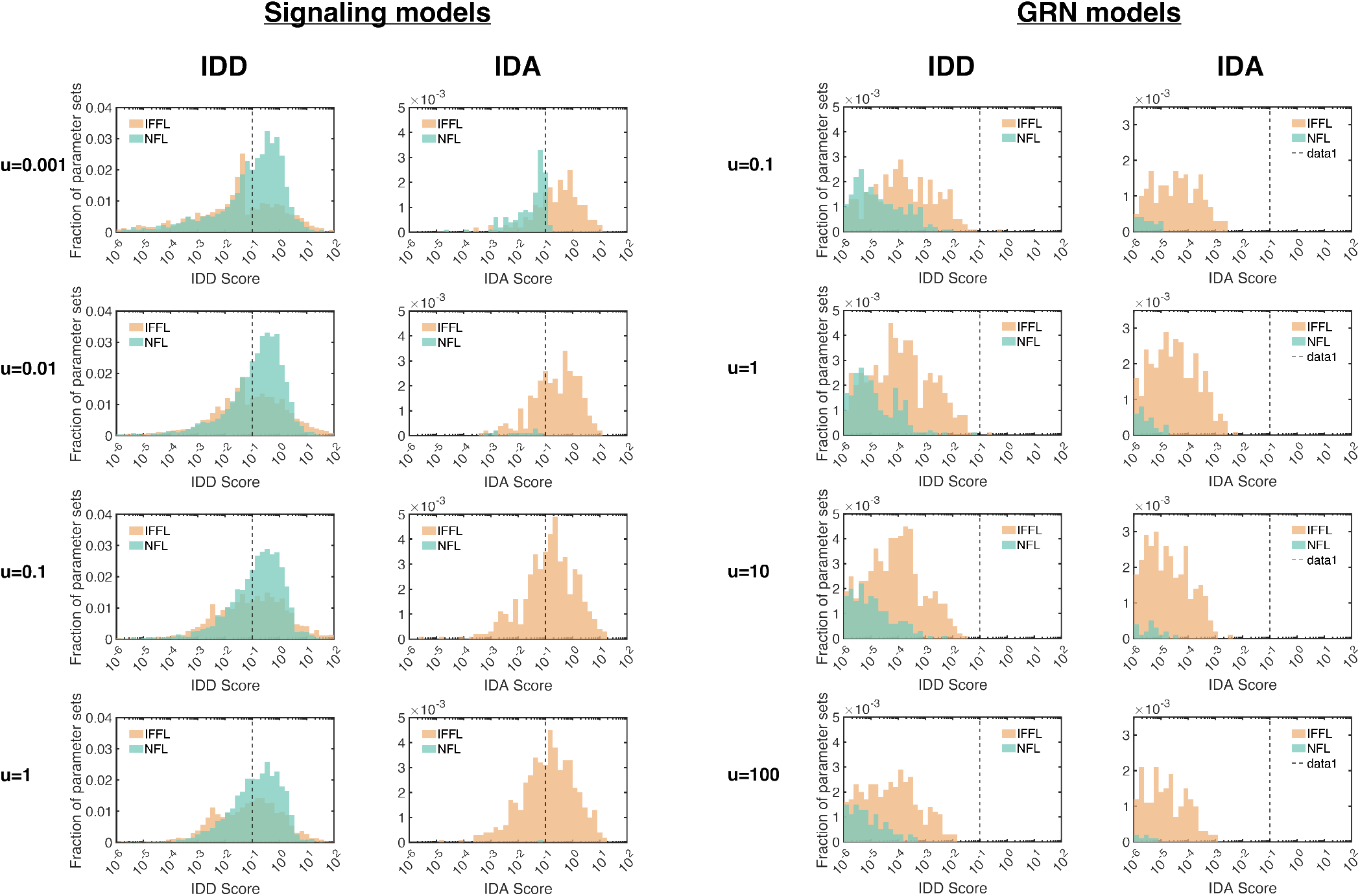
Histograms showing the fraction of parameter sets with positive IDD or IDA scores for the signaling and gene regulatory network (GRN) models of IFFL and NFL at different input amplitudes.

**Figure 17:**
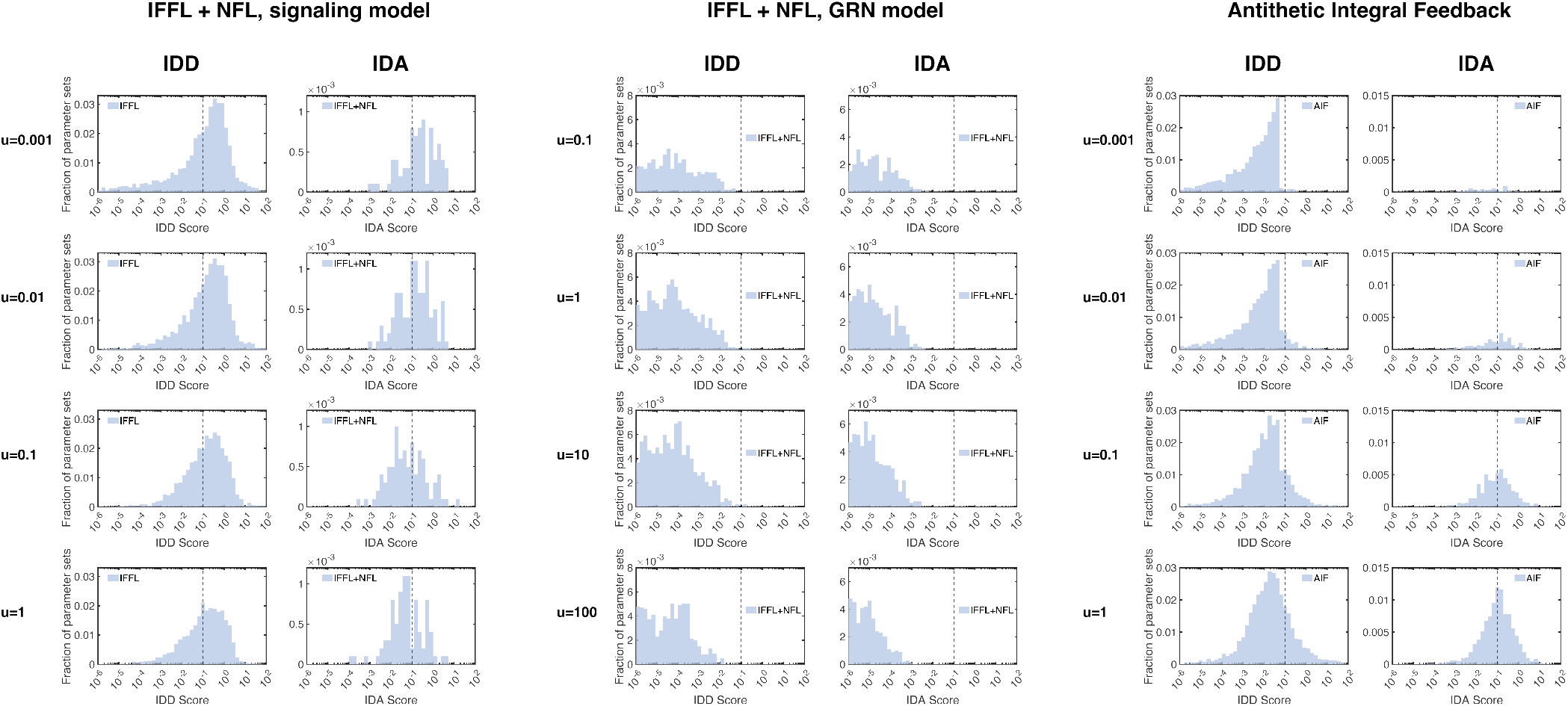
Histograms showing the fraction of parameter sets with positive IDD or IDA scores for the signaling and gene regulatory network (GRN) models of IFFL+NFL and the signaling model of the antithetic integral feedback circuit at different input amplitudes.

## Acknowledgments

E.N. and E.F acknowledge support from National Science Foundation through award NSF-CCF 2107483. F.B. and G.G. have been partly supported by the European Union through the Next Generation EU, Mission 4, Component 2, PRIN 2022 grant PRIDE (project number 2022LP77J4, CUP E53D2300072000); G.G. also acknowledges support from the European Union through the ERC INSPIRE grant (project number 101076926); views and opinions expressed are however those of the authors only and do not necessarily reflect those of the European Union, the European Research Council Executive Agency or the European Council; neither the European Union nor the granting authority can be held responsible for them. A.H. was supported by National Institutes of Health (R01AI173214). The authors thank all members of the labs for their support.

## Author contributions

Conceptualization, E.N., A.H., and E.F.; mathematical model development, E.N., F.B., G.G., and E.F.; numerical simulations, E.N.; mathematical analysis, F.B., G.G., and E.F.; experiments, E.N.; writing – original draft, E.N.; writing – review & editing, E.N., F.B., G.G, A.H., and E.F.; funding acquisition, E.F.; supervision, E.F.

## DECLARATION OF GENERATIVE AI AND AI-ASSISTED TECHNOLOGIES

During the preparation of this work, the authors used ChatGPT from OpenAI in order to refactor and write MATLAB code and to revise grammatical errors in the text. After using this tool, the authors reviewed and edited the content as needed and take full responsibility for the content of the publication.

## STAR METHODS

### Resource availability

#### Lead contact

Requests for further information and resources should be directed to and will be fulfilled by the lead contact, Elisa Franco.

#### Materials availability

This study did not generate new materials.

#### Data and code availability

- All the numerical simulation data used for the figures are publicly available as of the date of publication at https://github.com/enaka74/IDR.
- All original code has been deposited at https://github.com/enaka74/IDR and is publicly available as of the date of publication.
- All the raw data and the MATLAB scripts for analyzing the data have been deposited at https://zenodo.org/records/15385324.

### Method details

#### Mathematical models for the negative feedback loop (NFL) circuit

Temporarily denoting the concentration of species *Y* as 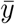, we introduce the simple NFL model as follows:

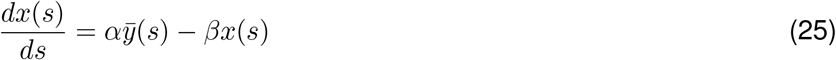

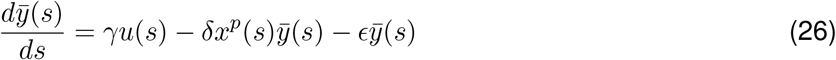

where *α* and *γ* are the activation rate constants for species *X* and *Y* respectively; *β* and *ϵ* are the self-degradation rate constants for species *X* and *Y* respectively; δ is the rate constant associated with inhibition of *Y* mediated by *X*, and *p* represents the order (i.e., cooperativity) of the reaction, while *s* is the time variable. To reduce the number of parameters, we rescale time in Equations 25 and 26 and obtain the new ODE model, which we will analyze in the following without loss of generality:

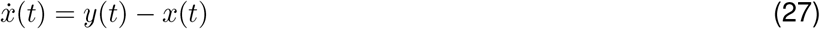

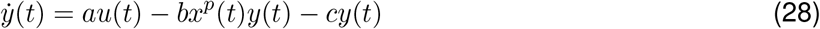

where *t* = *βs*, 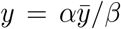, *a* = *γ/β, b* = δ*/α, c* = *ϵβ/α*, while 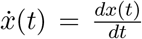 and 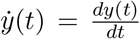. Note that variable *y* is rescaled here. Using Equations 27 and 28, we can evaluate the circuit dynamics with respect to *a, b, c*, and *p*, as done in the main text.

The signaling model of NFL is as follows:

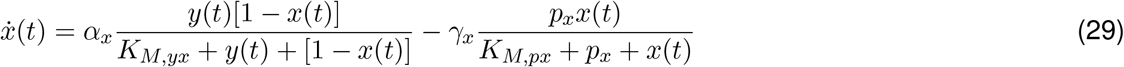

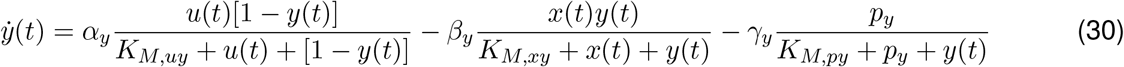

The gene regulatory model of NFL is as follows:

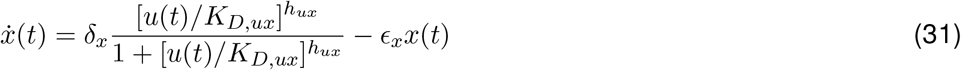

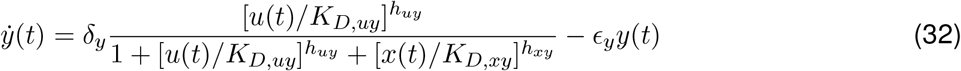

The parameters for each model are defined in the same way as those for IFFL.

#### Gene regulatory network model for the generalized incoherent feedforward loop (IFFL) circuit

We consider the following gene regulatory network model for the generalized incoherent feedforward loop:

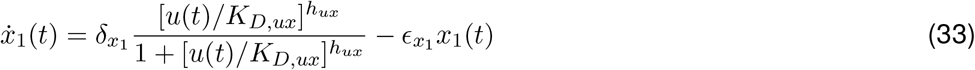

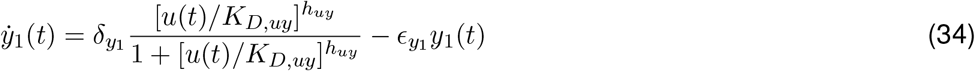

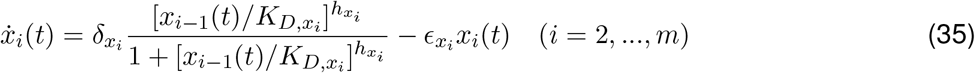

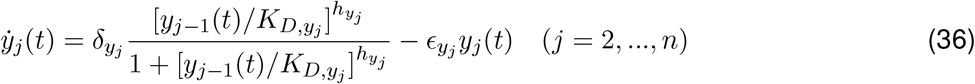

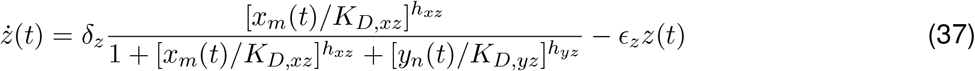

where δ is the activation rate and *ϵ* is the dilution/degradation rate for an expressed protein, while *K*_*D*_ is the dissociation constant of a transcription factor on its target promoter. The subscripts indicate the species in which a constant is involved; for example, 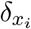 is the activation rate of *X* and 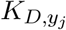 is the dissociation constant between the transcription factor expressed from the gene *Y*_*j−*1_ and the targeted promoter for gene *Y*_*j*_.

#### Parameter values for the detailed mathematical models

#### Quantification of the inverse properties

We introduce several mathematical formulae from Heinrich’s work^62^ to evaluate the transient dynamics of a signal. With the formulae, we can quantify temporal features of a signal of an arbitrary shape. Given the function *f* (*t*) of time, which can be the temporal signal obtained as the output response of a biological circuit, the formulae are as follows.

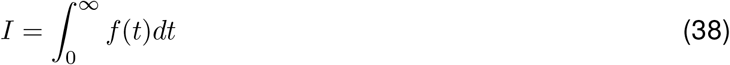

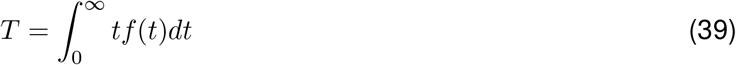

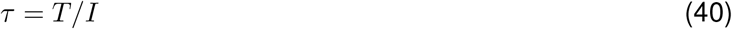

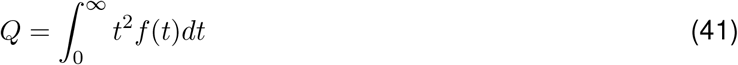

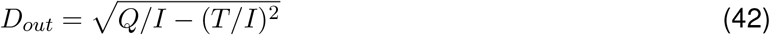

*I* is the integral of the signal, also referred to as the area under the curve or *AUC*, i.e. the integrated response. *T* is the time-weighted integrated response, while the *signaling time τ* is the average time required to activate the response. *Q* is the squared-time-weighted integrated response. The signal duration is denoted by *D*_*out*_. The quantities *τ* and *D*_*out*_ are analogous to the mean and standard deviation, respectively, of a continuous statistical distribution. Throughout the present study, we evaluate the signal duration of an output response by computing *D*_*out*_ as in Equation 42. It should be noted that *D*_*out*_ is undefined when *f* (*t*) = 0 due to division by zero. Specifically, *D*_*in*_ = 0 results in no response across all the models evaluated in the main text. Therefore, *D*_*out*_(*D*_*in*_ = 0) is not considered in this study.

Next we define and quantify the IDD property. We suppose *D*_*out*_(*D*_*in*_), namely the output signal duration as a function of the input signal duration, can have an arbitrary functional form, as shown in Fig. 18. The steps to compute the IDD score *S*_*IDD*_ are as follows: First, we locate all the local maxima, 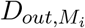, and local minima, 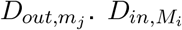 and 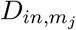 are the input pulse durations that give 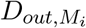 and 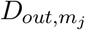 (i.e., 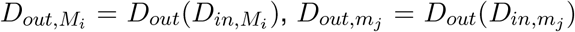)). Next we consider all combinations of 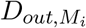 and 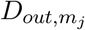. Finally, we determine the combination that gives the maximum value of 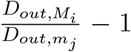, satisfying 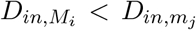, which guarantees the IDD property is maintained for all 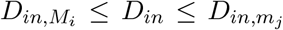. The following definition mathematically represents the IDD score *S*_*IDD*_ determined through the above steps:

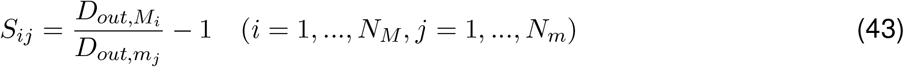

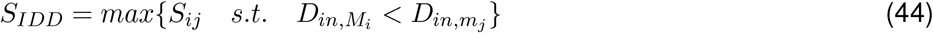

where *N*_*M*_ and *N*_*m*_ are the total number of maxima and minima, respectively. If *D*_*out*_(*D*_*in*_) is a non-decreasing function, then *S*_*IDD*_ is defined as zero. Because a constant function can be considered to have *D*_*out,M*_ = *D*_*out,m*_, which gives *S*_*IDD*_ = 0, this extension of the definition is reasonable. This expression can quantify the IDD property of an arbitrary shape of *D*_*out*_(*D*_*in*_) curve. Based on this definition, if a response has an interval of *D*_*in*_ values for which *D*_*out*_(*D*_*in*_) locally decreases, the response has a positive *S*_*IDD*_; otherwise, *S*_*IDD*_ = 0. An IDA score *S*_*IDA*_ is calculated in the same way, just considering function *I*(*D*_*in*_), namely *AUC*(*D*_*in*_).

**Figure 18:**
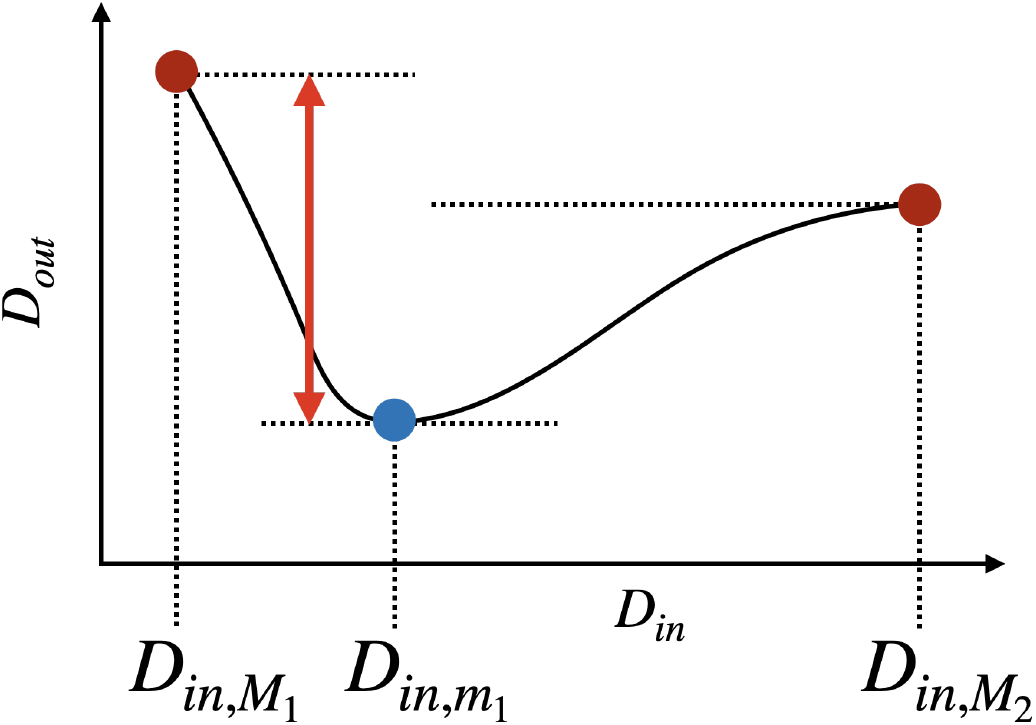
A schematic example of function *D*_*out*_(*D*_*in*_) and steps to find the 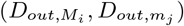 pair to compute the IDD score.

#### Computational simulations and random parameter sampling for the biological network models

All the numerical simulations were performed using MATLAB. The ode23s solver was used for the signaling models, and the ode15s solver was used for the GRN models. For random parameter sampling, each parameter (except for the Hill coefficient *h*) was sampled from the log-uniform distribution in the interval specified by the lower and upper limits listed in Table 5 using the MATLAB *rand* function. The Hill coefficient was sampled from a discrete uniform distribution over the integer values defined by the corresponding bounds in Table 5. The selected parameter ranges are biologically plausible and consistent in units, and are comparable to those used in previous modeling studies of biological circuits^46^. The input amplitude was varied among *u*_0_ = 0.001, 0.01, 0.1, 1 µM for the signaling models, and *u*_0_ = 0.1, 1, 10, 100 nM for the GRN models. The histograms selected for Figure 4 were generated using *u*_0_ = 0.01 µM for the signaling model and *u*_0_ = 1 nM for the GRN model. The remaining histograms are provided in the Supplemental Figure 16.

**Table 5:** Parameter ranges for the random parameter sampling.

| Parameter | Unit | Lower limit | Upper limit |
| --- | --- | --- | --- |
| $\alpha$ | $\text{min}^{-1}$ | $1 \times 10^{-1}$ | $1 \times 10^2$ |
| $\beta$ | $\text{min}^{-1}$ | $1 \times 10^{-1}$ | $1 \times 10^2$ |
| $\gamma$ | $\text{min}^{-1}$ | $1 \times 10^{-1}$ | $1 \times 10^2$ |
| $K_M$ | $\mu\text{M}$ | $1 \times 10^{-2}$ | $1 \times 10^2$ |
| $P$ | $\mu\text{M}$ | 0.1 | 1 |
| $\delta$ | $\text{nM hr}^{-1}$ | $1 \times 10^0$ | $1 \times 10^2$ |
| $\epsilon$ | $\text{hr}^{-1}$ | $1 \times 10^{-2}$ | $1 \times 10^0$ |
| $K_D$ | $\text{nM}$ | $1 \times 10^{-2}$ | $1 \times 10^2$ |
| $h$ | - | 1 | 4 |

**Table 6:** Concentration of genelets and buffer components in the experiments shown in Figure 3, Supplemental Figure 12 and 13.

| Component | Concentration |
| --- | --- |
| $G_{in}C_{out}$ | 25 nM<br>(10 nM, 25 nM, 50 nM in the experiments for the Supplemental Figure 13) |
| $G_{in}C_r$ | 5 nM<br>(2 nM, 5 nM, 10 nM in the experiments for the Supplemental Figure 13) |
| $dA_{in}$ | 150 nM |
| $dR_{in}$ | 250 nM |
| $G_rC_{out}$ | 5 nM |
| $dA_r$ | 50 nM |
| $dB_r$ | 125 nM |
| $G_{out}S$ | 25 nM |
| $G_{out}I_{out}$ | 1 nM |
| $dA_{out}$ | 100 nM |
| $dB_{out}$ | 37.5 nM |
| NTP | 7.5 mM each |
| $MgCl_2$ | 30 mM |
| Pyrophosphatase | 1.35 mU $\mu L^{-1}$ |
| Dithiothreitol | 5 mM |
| RNase H | 0.01 U $\mu L^{-1}$ |
| T7 Enzyme Mix | 2.9 U $\mu L^{-1}$ |

For the GRN models, additional time-scaling was conducted to improve the stability of the numerical simulations. To obtain a rough estimate of the overall time scale of the system with a given parameter set, we the following scaling factor:

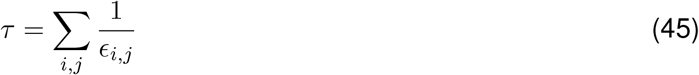

This scaling factor was applied to the time steps, absolute tolerance, and input durations.

For each sampled parameter set, simulations were performed over a range of input durations. For the signaling model, *D*_*in*_ was logarithmically swept over the interval [1, 100], and whereas for the GRN model, *D*_*in*_ was logarithmically swept over the interval [0.1, 10] ×*τ*. Note that, for the generalized IFFL networks with fixed parameter values and varying numbers of intermediate nodes, simulations were performed with *D*_*in*_ logarithmically swept over the interval [0.1, 10] in order to capture the range of durations over which the inversion property emerges.

#### Experimental Methods

##### Oligonucleotide sequences

All DNA was purchased from Integrated DNA Technologies (IDT, Coralville, Iowa, USA). The DNA sequences used for the genelet IFFL experiments are listed in Table 7. All sequences were adapted from genelets established by Schaffter et al.^21^. Specifically, G_in_C_out_, G_in_C_r_, G_r_R_out_, G_out_S, and G_out_I_out_ were derived from G8C1, G8C3, G3R1, G1S1, and G1I1, respectively, as reported in their study.

**Table 7:**
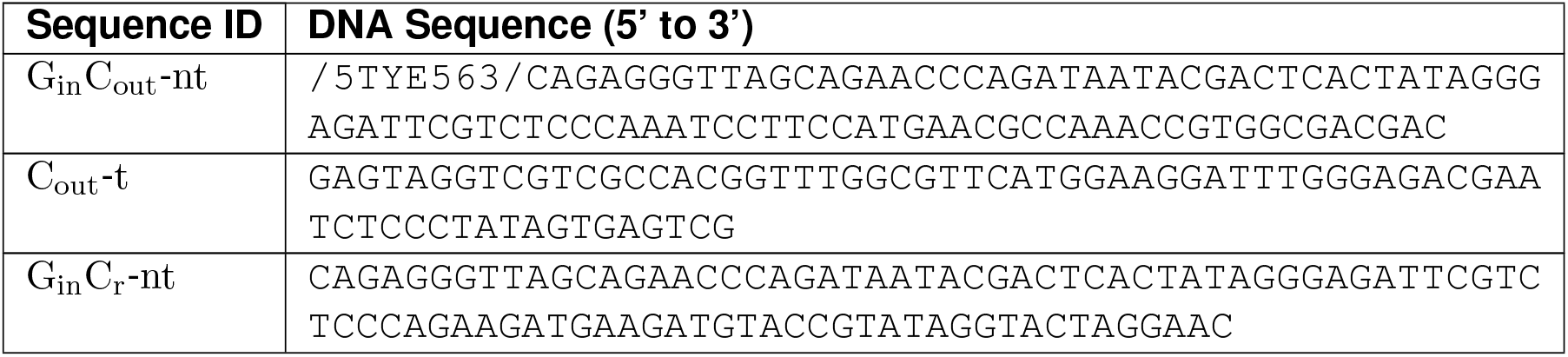
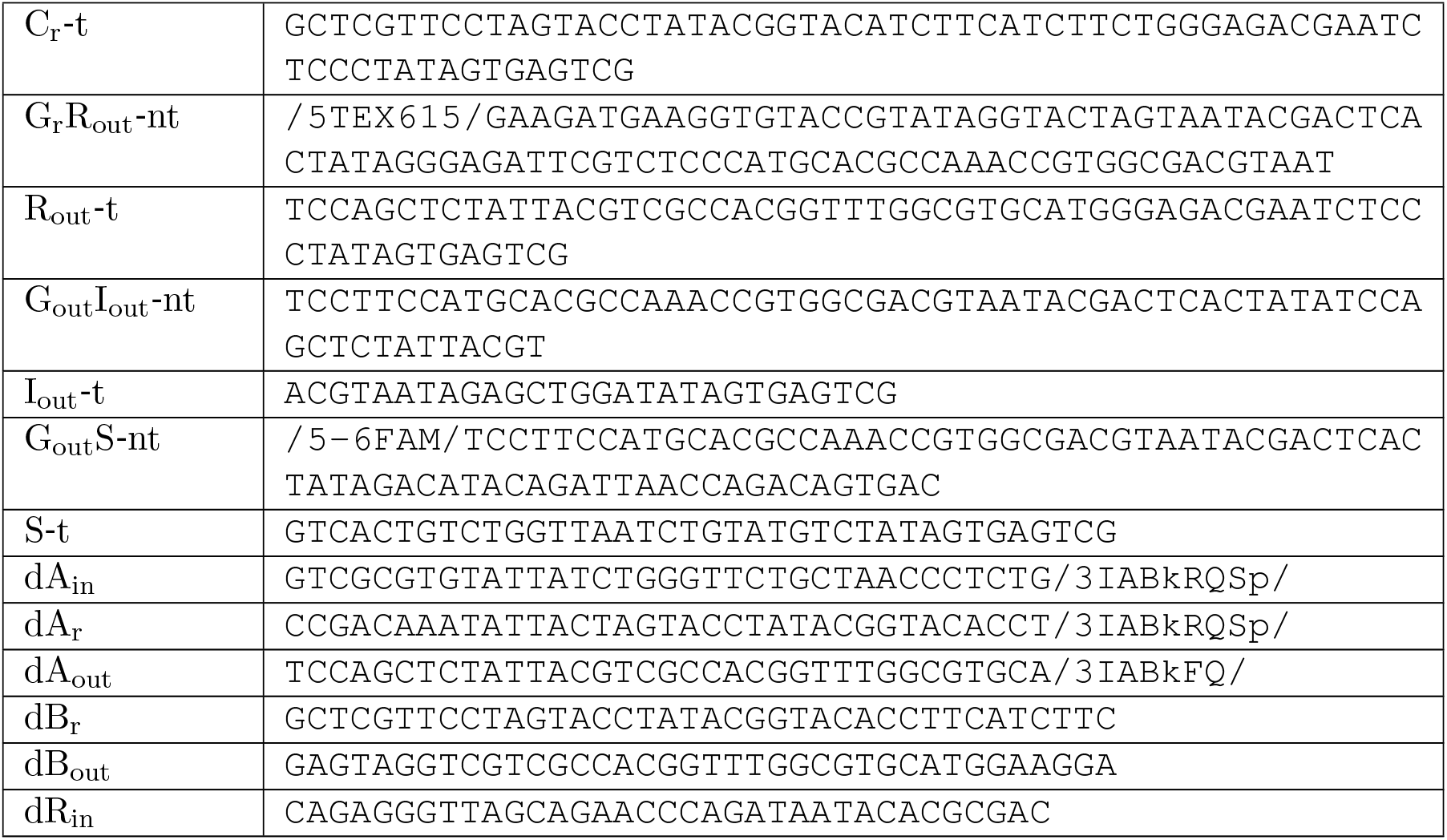
A list of the DNA sequences for the genelet IFFL experiments.

**Table 8:** Optimization bounds and initial values (placeholders to be filled manually).

| Parameter | Unit | Lower bound | Upper bound | Initial |
| --- | --- | --- | --- | --- |
| $k_{GA}$ | $M^{-1}s^{-1}$ | $1 \times 10^4$ | $1 \times 10^6$ | $1 \times 10^5$ |
| $k_{GAR}$ | $M^{-1}s^{-1}$ | $1 \times 10^3$ | $1 \times 10^5$ | $1 \times 10^4$ |
| $k_{GAR,d}$ | $M^{-1}s^{-1}$ | $5 \times 10^4$ | $5 \times 10^6$ | $5 \times 10^5$ |
| $k_{AR}$ | $M^{-1}s^{-1}$ | $1 \times 10^3$ | $1 \times 10^5$ | $1 \times 10^4$ |
| $k_{GBC}$ | $M^{-1}s^{-1}$ | $1 \times 10^2$ | $1 \times 10^4$ | $1 \times 10^3$ |
| $k_{GB}$ | $M^{-1}s^{-1}$ | $1 \times 10^4$ | $1 \times 10^6$ | $1 \times 10^5$ |
| $k_{GAB}$ | $M^{-1}s^{-1}$ | $1 \times 10^3$ | $1 \times 10^5$ | $1 \times 10^4$ |
| $k_{BC}$ | $M^{-1}s^{-1}$ | $1 \times 10^3$ | $1 \times 10^5$ | $1 \times 10^4$ |
| $k_{IR}$ | $M^{-1}s^{-1}$ | $1 \times 10^4$ | $1 \times 10^6$ | $1 \times 10^5$ |
| $k_{PR}$ | $s^{-1}$ | $1 \times 10^{-2}$ | $1 \times 10^0$ | $1 \times 10^{-1}$ |
| $k_{PC}$ | $s^{-1}$ | $1 \times 10^{-2}$ | $1 \times 10^0$ | $1 \times 10^{-1}$ |
| $k_{PI}$ | $s^{-1}$ | $1 \times 10^{-2}$ | $1 \times 10^0$ | $1 \times 10^{-1}$ |
| $k_{auto,input}$ | $s^{-1}$ | $1 \times 10^{-4}$ | $1 \times 10^{-2}$ | $1 \times 10^{-3}$ |
| $k_{auto,other}$ | $s^{-1}$ | $1 \times 10^{-5}$ | $1 \times 10^{-3}$ | $1 \times 10^{-4}$ |
| $k_{leak}$ | $s^{-1}$ | $2 \times 10^{-3}$ | $2 \times 10^{-1}$ | $2 \times 10^{-2}$ |
| $f_{P,exp4}$ | - | **0.9** | **1.1** | **1.0** |
| $k_{DR}$ | $s^{-1}$ | $1.5 \times 10^{-4}$ | $1.5 \times 10^{-2}$ | $1.5 \times 10^{-3}$ |
| $k_{DC}$ | $s^{-1}$ | $1.5 \times 10^{-4}$ | $1.5 \times 10^{-2}$ | $1.5 \times 10^{-3}$ |

##### Genelet preparation

Each genelet was annealed separately in 1X NEB RNAPol Reaction Buffer (New England Biolabs, Ipswich, MA, USA; catalog no. B9012SVIAL). Annealing protocol is as follows: heated to 90 ^*°*^C for 5 min and cooled to 20 ^*°*^C at a rate of 1 ^*°*^C min^*−*1^. The initially blocked genelets, G_r_R_out_, G_out_S, and G_out_I_out_, were annealed with a template strand, a non-template strand, and a DNA blocker at a 1.5-fold concentration. In contrast, the initially active genelets, G_in_C_out_ and G_in_C_r_, were annealed with only a template and a non-template strand.

#### Circuit preparation

Annealed IFFL genelets were combined in a single reaction containing NTPs (New England Biolabs; catalog no. N0450L), MgCl_2_, pyrophosphatase (New England Biolabs; catalog no. M2403L), and dithiothreitol (Promega, distributed by VWR, Radnor, PA, USA; catalog no. PAP1171) at the concentrations listed in Table 6.

#### Fluorescence spectroscopy

Fluorescence was measured by a spectrofluorometer (HORIBA Instruments Incorporated, Irvine, CA, USA; catalog no. FL3-11) or plate reader (), both of which are equipped with a temperature control system that maintained the sample temperature at 37 ^*°*^C.. The spectrofluormeter was used for the experiment shown in Figure 3E and F, and the plate reader was used for the experiments shown in Figure 3H.

For spectroflurometer measurements, samples were loaded into quartz cuvettes, and covered with 50 µL hexadecane to prevent evaporation. Cuvettes were placed in thespectrofluorometer. Fluorescence signals were recorded every two minutes at excitation/emission wavelengths of 495/520 nm for 6FAM dye, 549/565 nm for TYE563 dye, and 596/615 nm for TEX615 dye.

For plate reader measurements, samples were loaded into 384 Well Black Plate, Non-Treated Surface, Non-Sterile (Thermo Fisher Scientific, Waltham, MA, USA; catalog no. 262260), and covered with a VWR polyester film (Avantor, Radnor, PA, USA; catalog no. 60941-062) to prevent evaporation. Fluorescence signals were recorded every two minutes at excitation/emission wavelengths of 490/530 nm for 6FAM dye, 539/569 nm for TYE563 dye, and 590/621 nm for TEX615 dye.

##### Transcription reaction

Samples including all the IFFL genelets were loaded in the fluorimeter to collect baseline off fluorescence. After equilibration, a mixture of dA_in_, dA_r_, dA_out_, dB_r_, and dB_out_ was added. The amounts of dB_r_ and dB_out_ were adjusted such that the total concentrations of [dB_r_] and [dB_out_], including the contributions from dB_r_ and dB_in_ added during annealing, matched the values listed in Table 6. At this point, only G_in_C_out_ and G_in_C_r_ are active because the remaining genelets are in a blocked state. After the fluorescence signals stabilized, RNase H (Thermo Fisher Scientific, Waltham, MA, USA; catalog no. EN0202) and T7 Enzyme mix from MEGAshortscript T7 Transcription Kit (Invitrogen, Thermo Fisher Scientific; catalog no. AM1354) were added. The final concentrations correspond to those in Table 6 with a total reaction volume of 60 µL for fluorometer measurement and 20 µL for plate reader measurement. After a predetermined time interval, the DNA repressor dR_in_ was introduced to repress the input genelets (G_in_C_out_ and G_in_C_r_), resulting in the pulsatile activation of the input genelets.

##### Fluorescence normalization

The active level of a fluorescently tagged genelet was calculated based on the maximum fluorescence *F*_*M*_ (fluorescence provided by fluorescently tagged genelets only) and the minimum fluorescence *F*_*m*_ (fluorescence from a fully-quenched genelet). Note that both *F*_*M*_ and *F*_*m*_ are time-course trajectories obtained from control experiments, so hereinafter we denote *F*_*M*_ (*t*) and *F*_*m*_(*t*) for clarity. First, in control samples, two trajectories were obtained: *F*_*ctr,M*_ (*t*) a trajectory reflecting the maximum fluorescence value, and *F*_*ctr,m*_(*t*), a trajectory reflecting the minimum fluorescence value. *F*_*ctr,M*_ was obtained from a sample to which no activators were added, and *F*_*ctr,m*_ was obtained by adding excess amount of activators to a sample that has no blockers (Figure 19). Therefore, *F*_*ctr,M*_ and *F*_*ctr,m*_ are supposed to show the maximum and minimum fluorescence signals corresponding to 0 % activation and 0 % activation, respectively. However, *F*_*ctr,M*_ and *F*_*ctr,m*_ have to be adjusted based on sample-to-sample intensity variation. Therefore, *F*_*M*_ and *F*_*m*_ were calculated by multiplying a scaling factor *f*_*M*_ and *f*_*m*_. *f*_*M*_ and *f*_*M*_ is calculated based on a relative fluorescent intensity at a specified time point, *t*_*ref*_. Thus

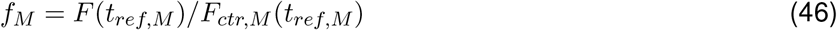

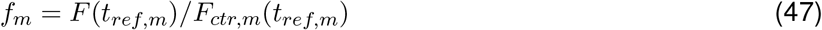

Thus, adjusted trajectories of the maximum and minimum fluorescence are:

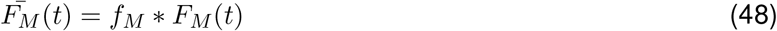

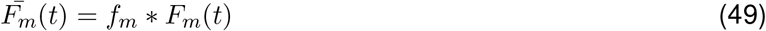

**Figure 19:**
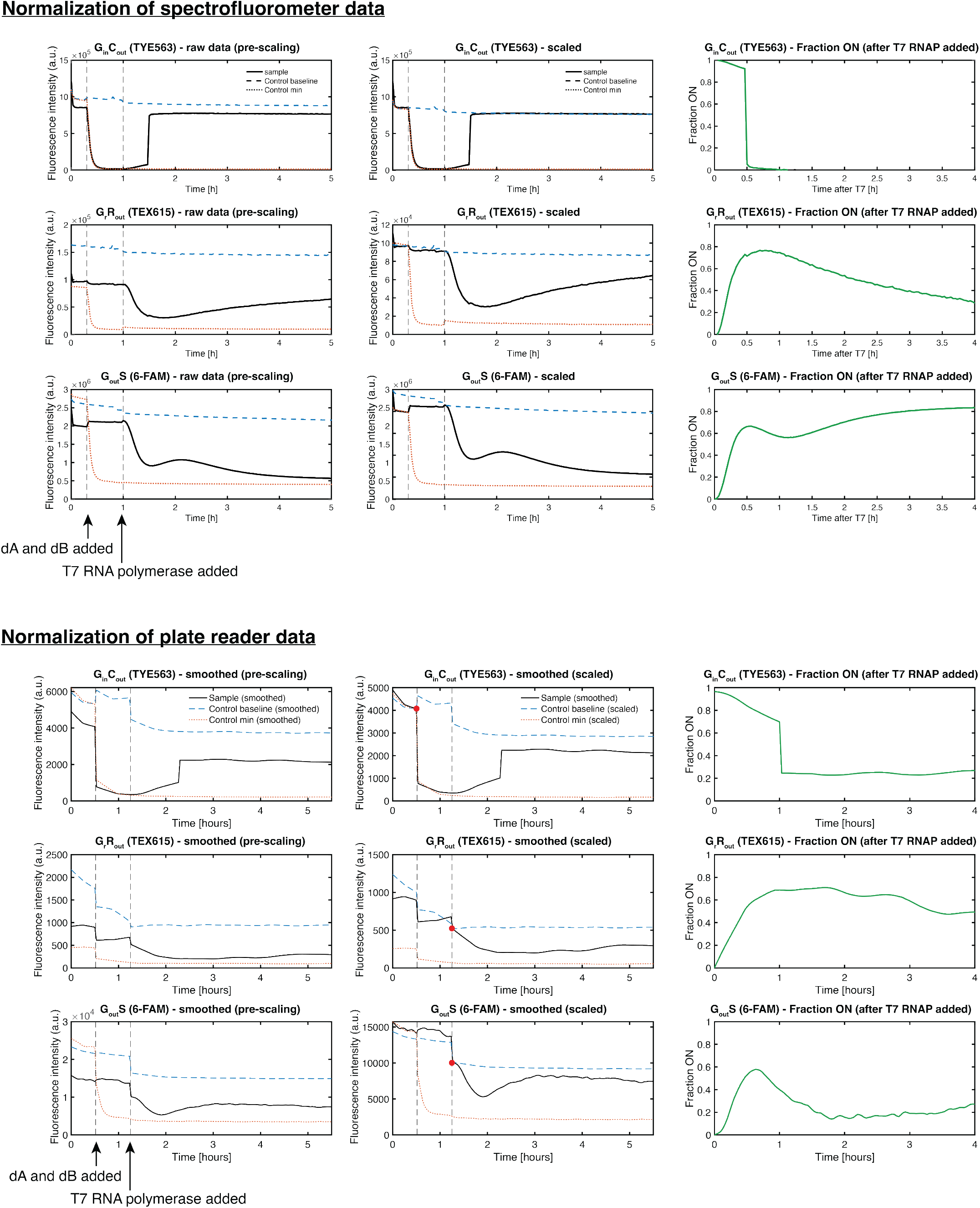
Normalization examples for sperctrofluorometer dataset (top) and plate reader dataset (bottom). Each dataset has fluorescence data at three wavelength (corresponding to each row). For the spectrofluorometer dataset, the left column shows raw data, the central column shows scaled data, and the right column shows normalized data. For the plate reader data, the left column shows smoothed data before scaling, the central column shows smoothed and scaled data, and the right column shows normalized smoothed data.

Finally, the active level of a genelet at time *t*, [*G*]_*on*_(*t*), is calculated as follows:

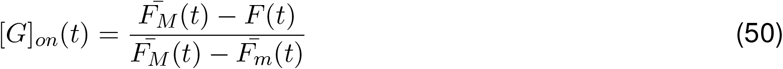

For spectrofluorometer data, fluorescence signal was also scaled based on volume change every time additional reagents (e.g., DNA activators and enzymes) are added, as the spectrofluorometer measurement is more sensitive to dilution than the plate reader measurement.

The whole normalization procedure is visualized in Figure 19 with example data. The spectrofluorometer has much larger signal-to-noise ratio, enabling dana analysis without any smoothing. Furthermore, the fluorometer data is showing more reasonable normalization results: for instance, *G*_*in*_*C*_*r*_ active fraction shows complete shut off after DNA repressor was added.

#### Parameter fitting of the genelet model

Parameter fitting of the genelet IFFL model was performed in MATLAB using a combination of the genetic algorithm function `ga` and the constrained nonlinear optimization function `fmincon`.

In the first fitting round, all model parameters were fitted to the time-series data of the output genelet *G*_*out*_*S* in response to input pulse durations of 10, 20, 30, 40, 60, 240 min (Fig. 3E) using the genetic algorithm function ga. At this stage, all kinetic parameters were constrained to be identical across nodes; for example, the *G*_*in*_, *G*_*r*_, and *G*_*out*_ domains shared a common value of *k*_*GA*_. Lower and upper bounds for each parameter were set to one order of magnitude above and below the corresponding initial value. The fitness function for this first fitting round was defined as

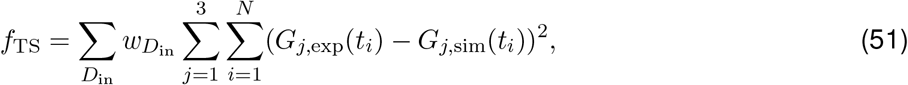

where *D*_in_ denotes the input pulse duration, *t*_*i*_ denotes the *i*-th experimental sampling time point for a given *D*_in_, and *N* is the number of sampled time points under that condition. The index *j* corresponds to the genelet species, with *j* = 1, 2, 3 representing *G*_*in*_, *G*_*r*_, and *G*_*out*_, respectively. *G*_*j*,exp_ and *G*_*j*,sim_ denote the experimentally measured and simulated active fractions of genelet *j*. The weights 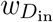 and *w*_*j*_ control the relative contribution of each input duration and genelet species to the overall objective function.

In the second fitting round, the parameters *k*_*GA*_ and *k*_*GAR*_ associated with the *G*_*out*_*S* node, as well as *k*_*pc*_, *k*_*pr*_, *k*_*dc*_, and *k*_*dr*_ for all nodes, were independently fitted, while all other parameters were held fixed at the values obtained from the first fitting step. The parameters were simultaneously fitted to the time-series data shown in Fig. 3E and to experimentally measured AUC values obtained by varying [*G*_*r*_*R*_*out*_]_tot_ (Experiment (i) in Fig. 3H), again using the MATLAB genetic algorithm function `ga`. The resulting parameter set was subsequently refined using the MATLAB function `fmincon`. The fitness function used in this round was defined as

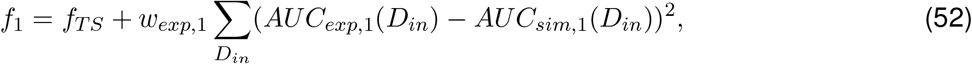

where *AUC*_*exp*,1_ and *AUC*_*sim*,1_ denote the experimentally measured and simulated AUC values, respectively, for Experiment (i) in Fig. 3H.

In the third and fourth fitting rounds, binding rate constants associated with different toehold lengths were fitted separately to Experiments (ii) and (iii) in Fig. 3 using the ga function. In the third step, the parameters *k*_*ar*_ and *k*_*gar*_ corresponding to 6-nt and 4-nt *dA*_*out*_ toeholds were fitted, while in the fourth step, the parameters *k*_*bc*_ and *k*_*gbc*_ corresponding to 6-nt and 4-nt *dB*_*out*_ toeholds were fitted. In both steps, binding rate constants were expressed as scaling factors relative to the values for the nominal toehold length (8 nt) and were constrained to lie within the range 10^*−*3^ to 1. The fitness functions for the third and fourth fitting rounds were defined as

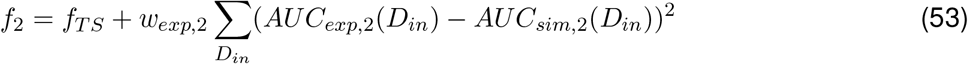

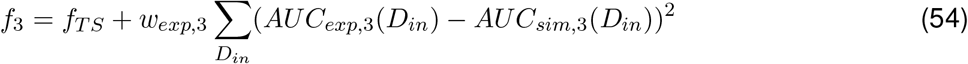

In the fifth fitting round, a scaling factor only for the experiment with varying *G*_*in*_*C*_*out*_ and *G*_*in*_*C*_*out*_ concentrations (Fig. 3F) (right) was fitted. This scaling factor is used only for the experiment, as the experiment was performed with a different batch of T7 RNA polymerase, which is assumed to uniformly affects all the RNA production rate constants. The fitness function for the fifth fitting round was defined as

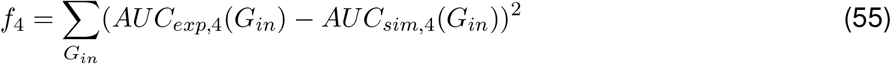

Through these fitting rounds, the parameter values listed in Table 9, 10, 11, and 12 were obtained.

**Table 9:** Common parameters shared across all nodes (values only).

| Parameter | Unit | Value |
| --- | --- | --- |
| $k_{GA}$ | $M^{-1}s^{-1}$ | $9.12 \times 10^4$ |
| $k_{GAR,d}$ | $M^{-1}s^{-1}$ | $4.64 \times 10^6$ |
| $k_{GB}$ | $M^{-1}s^{-1}$ | $1.71 \times 10^4$ |
| $k_{GAB}$ | $M^{-1}s^{-1}$ | $1.27 \times 10^4$ |
| $k_{IR}$ | $M^{-1}s^{-1}$ | $7.02 \times 10^4$ |
| $k_{auto,input}$ | $s^{-1}$ | $1.11 \times 10^{-3}$ |
| $k_{auto,other}$ | $s^{-1}$ | $5.42 \times 10^{-4}$ |
| $k_{leak}$ | $s^{-1}$ | $4.35 \times 10^{-3}$ |

**Table 10:** *m*-dependent parameters (*m* = 5), values only.

| Parameter | Unit | $G_{in}C_{out}$ | $G_{in}C_r$ | $G_rR_{out}$ | $G_{out}I_{out}$ | $G_{out}S$ |
| --- | --- | --- | --- | --- | --- | --- |
| $k_{PR}$ | $s^{-1}$ | - | - | $1.68 \times 10^{-2}$ | - | - |
| $k_{PC}$ | $s^{-1}$ | $3.96 \times 10^{-2}$ | $8.06 \times 10^{-2}$ | - | - | - |
| $k_{PI}$ | $s^{-1}$ | - | - | - | $2.97 \times 10^{-2}$ | - |
| $f_{P,exp4}$ | - | 1.10 | 1.10 | 1.10 | 1.10 | 1.10 |

**Table 11:** *n*-dependent parameters (*n* = 3), values only.

| Parameter | Unit | $G_{in}$ | $G_r$ | $G_{out}$ |
| --- | --- | --- | --- | --- |
| $k_{DR}$ | $s^{-1}$ | $3.52 \times 10^{-3}$ | $1.86 \times 10^{-3}$ | $9.02 \times 10^{-4}$ |
| $k_{DC}$ | $s^{-1}$ | $3.52 \times 10^{-3}$ | $1.86 \times 10^{-3}$ | $9.02 \times 10^{-4}$ |
| $k_{GAR}$ | $M^{-1}s^{-1}$ | $1.34 \times 10^4$ | $1.34 \times 10^4$ | $3.67 \times 10^5$ |
| $k_{AR}$ | $M^{-1}s^{-1}$ | $1.16 \times 10^4$ | $1.16 \times 10^4$ | $6.93 \times 10^4$ |
| $k_{GBC}$ | $M^{-1}s^{-1}$ | $4.12 \times 10^3$ | $4.12 \times 10^3$ | $4.12 \times 10^3$ |
| $k_{BC}$ | $M^{-1}s^{-1}$ | $4.08 \times 10^3$ | $4.08 \times 10^3$ | $4.08 \times 10^3$ |

**Table 12:** Tuned parameters for Exp2 and Exp3.

| Parameter | Unit | Value | Scale range |
| --- | --- | --- | --- |
| $k_{AR}$ (8 nt) | $M^{-1}s^{-1}$ | $6.93 \times 10^4$ | [0.001, 1] |
| $k_{AR}$ (6 nt) | $M^{-1}s^{-1}$ | $5.48 \times 10^4$ | [0.001, 1] |
| $k_{AR}$ (4 nt) | $M^{-1}s^{-1}$ | $4.48 \times 10^3$ | [0.001, 1] |
| $k_{GAR}$ (8 nt) | $M^{-1}s^{-1}$ | $3.67 \times 10^5$ | [0.001, 1] |
| $k_{GAR}$ (6 nt) | $M^{-1}s^{-1}$ | $4.98 \times 10^4$ | [0.001, 1] |
| $k_{GAR}$ (4 nt) | $M^{-1}s^{-1}$ | $6.92 \times 10^3$ | [0.001, 1] |
| $k_{BC}$ (8 nt) | $M^{-1}s^{-1}$ | $4.08 \times 10^3$ | [0.001, 1] |
| $k_{BC}$ (6 nt) | $M^{-1}s^{-1}$ | $4.09 \times 10^0$ | [0.001, 1] |
| $k_{BC}$ (4 nt) | $M^{-1}s^{-1}$ | $4.38 \times 10^0$ | [0.001, 1] |
| $k_{GBC}$ (8 nt) | $M^{-1}s^{-1}$ | $4.12 \times 10^3$ | [0.001, 1] |
| $k_{GBC}$ (6 nt) | $M^{-1}s^{-1}$ | $1.18 \times 10^2$ | [0.001, 1] |
| $k_{GBC}$ (4 nt) | $M^{-1}s^{-1}$ | $4.23 \times 10^0$ | [0.001, 1] |

